# Membrane potential resonance in non-oscillatory neurons interacts with synaptic connectivity to produce network oscillations

**DOI:** 10.1101/394650

**Authors:** Andrea Bel, Horacio G. Rotstein

**Author notes:** Corresponding Investigator, CONICET, Argentina.

## Abstract

Several neuron types have been shown to exhibit (subthreshold) membrane potential resonance (MPR), defined as the occurrence of a peak in their voltage amplitude response to oscillatory input currents at a preferred (resonant) frequency. MPR has been investigated both experimentally and theoretically. However, whether MPR is simply an epiphenomenon or it plays a functional role for the generation of neuronal network oscillations and how the latent time scales present in individual, non-oscillatory cells affect the properties of the oscillatory networks in which they are embedded are open questions. We address these issues by investigating a minimal network model consisting of (i) a non-oscillatory linear resonator (band-pass filter) with 2D dynamics, (ii) a passive cell (low-pass filter) with 1D linear dynamics, and (iii) nonlinear graded synaptic connections (excitatory or inhibitory) with instantaneous dynamics. We demonstrate that (i) the network oscillations crucially depend on the presence of MPR in the resonator, (ii) they are amplified by the network connectivity, (iii) they develop relaxation oscillations for high enough levels of mutual inhibition/excitation, and the network frequency monotonically depends on the resonators resonant frequency. We explain these phenomena using a reduced adapted version of the classical phase-plane analysis that helps uncovering the type of effective network nonlinearities that contribute to the generation of network oscillations. Our results have direct implications for network models of firing rate type and other biological oscillatory networks (e.g, biochemical, genetic).

**Author Summary:** Biological oscillations are ubiquitous in living systems and underlie fundamental processes in healthy and diseased individuals. Understanding how the intrinsic oscillatory properties of the participating nodes interact with the network connectivity is key for the mechanistic description of biological network oscillations. In several cases these intrinsic oscillatory properties are hidden and emerge only in the presence of external oscillatory inputs in the form of preferred amplitude responses to these inputs. This phenomenon is referred to as resonance and may occur in systems that do not exhibit intrinsic oscillations. Resonance has been primarily measured in neuronal systems, but their role in the generation of neuronal network oscillations remains largely an open question. We have identified a minimal network model consisting of a resonator (a node that exhibits resonance, but not intrinsic oscillations), a low-pass filter (no resonance and no intrinsic oscillations) and nonlinear connectivity with no dynamics. This network is able to produce oscillations, even in the absence of intrinsic oscillatory components. These oscillations crucially depend on the presence of the resonator. Moreover, the resonant frequency, a dynamic property of the interaction between the resonator and oscillatory inputs, controls the network frequency in a monotonic fashion. The results of our study have implications for the generation of biological network oscillations in larger neuronal systems and other biological networks.

## 1 Introduction

Neuronal network oscillations emerge from the cooperative activity of the participating neurons and the network connectivity and involve the interplay of the nonlinearities and time scales present in the ionic and synaptic currents. In some cases, the network time scales directly reflect the time scales of the individual neurons. This class includes the networks that synchronize the oscillatory activity of the individual neurons where the frequency of both (network and individual neurons) belong to the same (narrow) frequency band. In other cases, the oscillatory time scales are latent (or hidden) at the individual neuron level and become apparent at the network level. The oscillatory networks of non-oscillatory neurons we investigate in this paper belong to this class. We focus on the situations where at least one of the participating cells exhibits (subthreshold) membrane potential resonance (MPR), a peak in the voltage amplitude response to oscillatory input currents at a preferred (resonant) frequency [1–4]. Because the individual cells are non-oscillatory, the resonant frequency is an oscillatory latent time scale.

The mechanisms of generation of sustained (limit cycle) oscillations in single neurons are reasonably well understood [5–8]. They require the interplay of negative and positive feedback effects provided by the ionic current gating variables or related processes. Resonant ionic processes (e.g., hyperpolarization-activated mixed-cation *I*_*h*_ current, M-type slow-potassium current *I*_*Ks*_ and T-type calcium inactivation *I*_*CaT*_) oppose changes in voltage, while amplifying ionic processes (e.g., persistent sodium current *I*_*Nap*_, T-type calcium activation) favor these changes.

There is a hierarchy of dynamic oscillatory phenomena that requires the presence of a resonant process and whose degree complexity increases with the levels of the amplifying current [1,9] in system where sustained oscillations (subthreshold or spikes) are generated by Hopf bifurcation mechanisms [5–7]. At the bottom of this hierarchy are the overshoot type of responses to square-pulse perturbations (Fig. 1, green curves) in neurons that exhibit MPR [1–4], but not subthreshold oscillations (STOs). We refer to them as resonators. For higher amplification levels the neuron may display damped subthreshold oscillations (Fig. 1, red curves). In these two cases the underlying systems may be quasi-linear in large enough vicinities of the resting potential (fixed-point) [9]. At the top of the hierarchy are the limit cycle oscillations (Fig. 1, blue curves) that require high enough amplification levels for the development of the nonlinearities necessary for the existence of limit cycles [9]. If these limit cycles represent STOs, additional amplification levels can produce spikes or depolarization block. Examples of models exhibiting this type of behavior are the Morris-Lecar model [10] and the *I*_*h*_ + *I*_*Nap*_ or *I*_*Ks*_ + *I*_*Nap*_ models studied in [9] (see also [11]).

**Figure 1:**
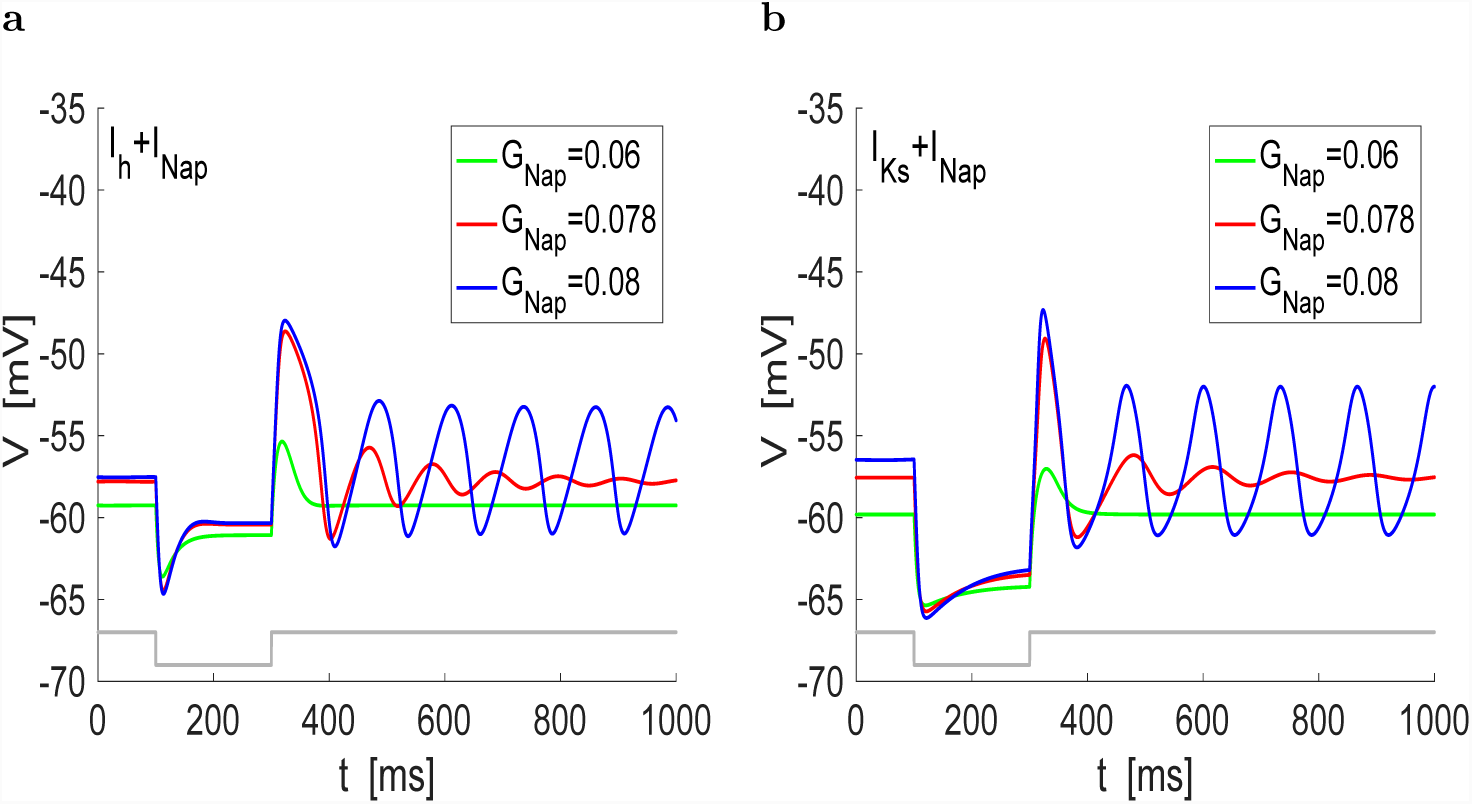
Response of *I*_*h*_+*I*_*Nap*_ and *I*_*Ks*_+*I*_*Nap*_ models to negative square pulses of current: representative dynamic scenarios. **a.** *I*_*h*_+*I*_*Nap*_ model. It includes three ionic currents: hyperpolarization-activated (h-), persistent sodium and leak (see Section 2.4 in Methods). **b.** *I*_*Ks*_+*I*_*Nap*_ model. It includes three ionic currents: M-type slow potassium, persistent sodium and leak (see Section 2.4 Methods). Both *I*_*h*_ and *I*_*Ks*_ are resonant and *I*_*Nap*_ is amplifying. Increasing the levels of *I*_*Nap*_ causes a transition from overshoot responses (green) to damped oscillations (red) to persistent (limit cycle) oscillations (blue) in both models. The gray curve is a caricature of the square wave input deflected from zero with amplitude 1. We used the following parameter values: *C* = 1, *E*_*Na*_ = 42, *E*_*L*_ = −75, *E*_*h*_ = −26, *G*_*L*_ = 0.3, *G*_*h*_ = 1.5, *I*_*app*_ = 0.55, *V*_*hlf,p*_ = −54.7, *V*_*slp,p*_ = 4.4, *V*_*hlf,q*_ = −80.2 and *V*_*slp,q*_ = 7.2 (*I*_*h*_+*I*_*Nap*_ model) and *C* = 1, *E*_*Na*_ = 42, *E*_*L*_ = −75, *E*_*Ks*_ = −96, *G*_*L*_ = 0.3, *G*_*Ks*_ = 1.5 and *I*_*app*_ = 4, *V*_*hlf,p*_ = −54.7, *V*_*slp,p*_ = 4.4, *V*_*hlf,q*_ = −28, *V*_*slp,q*_ = 8 (*I*_*Ks*_+*I*_*Nap*_ model).

MPR has been investigated in many neuron types both experimentally and theoretically [1–4, 12– 56]. However, in contrast to single cell intrinsic oscillations, the role that cellular MPR plays in network oscillations is not well understood. Only a few studies have addressed these issues in networks having neurons that exhibit MPR [41, 57–63] or have resonant gating variables [64–66]. To our knowledge, no study to date has examined the detailed mechanisms of generation of oscillations in networks of non-oscillatory resonators and how the network oscillations reflect the latent time scale provided by the resonant frequency.

In this paper we seek to understand how the resonant properties of individual nodes interact with the network connectivity to produce oscillations in reciprocally connected networks. We reasoned that if oscillations are to be generated in networks where the participating neurons only provide the resonant properties, then the amplification effects should result from the network connectivity. According to this hypothesis, oscillations will be generated in self-excited, but not self-inhibited resonators and in two-cell networks of mutually inhibited or mutually excited cells that include one resonator. Moreover, the resonant frequency of the individual resonators should control, or at least have a direct effect, on the network frequency. Along these lines, self-excited passive cells and mutually inhibited passive cells should not be able to oscillate. Analogously to single cell oscillations, the mechanism of generation of network oscillations should involve a Hopf bifurcation and the dynamic hierarchy described above. Some of these patterns have been observed for similar systems [67] and for network models using the Wilson-Cowan formalism [5, 68, 69]. However, the role that the filtering properties of the individual nodes (preferred frequency responses to oscillatory inputs) play in the generation of network oscillations has not been investigated.

We test these ideas using the simplest types of oscillatory networks with non-oscillatory neurons, consisting of a linear resonator reciprocally connected to a linear low-pass filter or another resonator with instantaneous graded synapses. More specifically, we use linear (linearized conductance-based) models for the individual neurons to isolate the resonant effects from the nonlinear amplifications that lead to sustained oscillations. In this way, we eliminate the nonlinear amplification from the dynamics of the individual neurons in order to asses the effects of subthreshold resonance on the network oscillations. These linearized models capture the quasi-linear dynamics of models having the passive currents and *I*_*h*_ or *I*_*Ks*_, but no amplifying currents (e.g., *I*_*Nap*_) [9]. They also capture the linearized dynamics of firing rate models of Wilson-Cowan type [69] with adaptation [70–72]. We primarily focus on parameter regimes where the individual neurons are resonators (produce overshoot responses to square pulses of current, without damped oscillations). We use graded synapses because of both the dynamic properties of linearized models and the range of voltage in which they operate and because it is the type of nonlinearities used in firing rate models. They are assumed to be instantaneously fast and have no dynamics [64, 65, 67, 70–75].

The questions we ask in this paper are conceptually different from the issues addressed in previous studies [41,64,65,67,73] and aim to conceptually address the mechanisms by which neuronal frequency filters interact within a network. This has implications not only for the understanding of neuronal oscillations, but also for the understanding how frequency-dependent information is communicated across neurons and networks and the phenomenon of network resonance [57, 76].

This paper is motivated by previous work on neuronal resonance in individual neurons [3, 4, 11, 40, 41, 57], and therefore we formulate the problem and the results in terms of neuronal systems, using the typical terminology and notation in the field of neuroscience. However, the issues addressed in this paper are more general and have direct implications to biological oscillatory networks in other fields (e.g., biochemical, genetic).

## 2 Methods

### 2.1 Networks of linearized cells with graded synapses

We used linearized biophysical (conductance-based) models for the individual cells and (nonlinear) graded synaptic connections. The linearization process for conductance-based models for single cells has been previously described in [2, 3]. We refer the reader to these references for details.

The dynamics of a two-cell network are described by

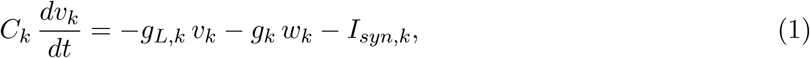

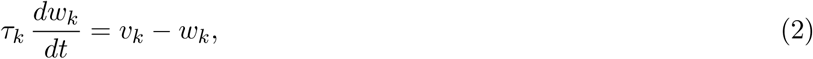

for *k* = 1, 2. In eqs. (1)-(2) *t* is time, *v*_*k*_ is the voltage (mV) referred to the voltage coordinate of the fixed-point (equilibrium potential) 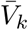, *w*_*k*_ is the gating variable referred to the gating variable coordinate of the fixed-point 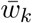 and normalized by the derivative of the corresponding activation curve, *C*_*k*_ is the capacitance, *g*_*L,k*_ is the linearized leak maximal conductance, *g*_*k*_ is the ionic current linearized conductance, *τ*_*k*_ is the linearized time constant and *I*_*syn,k*_ is the graded synaptic current from the other neuron in the network and given by

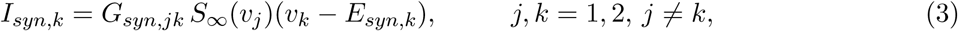

where *G*_*syn,k*_ is the maximal synaptic conductance, *E*_*syn,k*_ is the synaptic reversal potential referred to 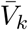 and

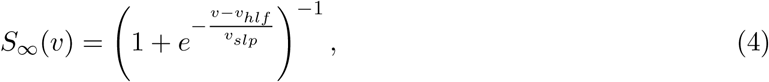

where the half-activation point *v*_*hlf*_ is also referred to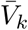.

We use the following units: mV for voltage, ms for time, *μ*F/cm^2^ for capacitance, *μ*A/cm^2^ for current and mS/cm^2^ for the maximal c1onductances. Unless stated otherwise, we used the following parameter values: *C* = 1, *V*_*hlf*_ = 0, *V*_*slp*_ = 1, *E*_*in*_ = −20, *E*_*ex*_ = 60.

Note that the heterogeneity due to different values of the DC current *I*_*app,k*_ and other biophysical parameters in the original conductance-based model is translated into the reversal potentials *E*_*syn,k*_ and the functions *S*_*k,∞*_(*v*) through the fixed-point 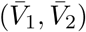. Specifically, if *E*_*syn*_ and *V*_*hlf*_ are the synaptic reversal potential and synaptic half-activation point of the original (not rescaled) model, then 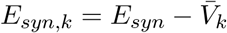 and 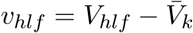.

### 2.2 Phase-space diagrams: nullclines and hyper-nullclines

#### 2.2.1 Graded network of two 1D passive cells: nullclines

These are 2D networks consisting of system (1) with *g*_*k*_ = 0 (*k* = 1, 2). The *v*_1_- and *v*_2_-nullclines are given by

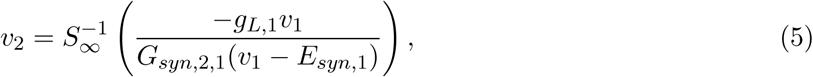

and

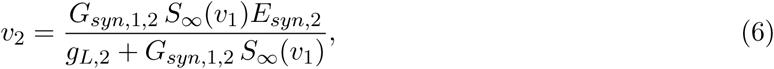

respectively, where

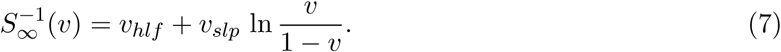

#### 2.2.2 Self-connected graded networks of a 2D resonant cell: nullclines

These networks are given by system (1)-(2) where *v*_2_ is substituted by *v*_1_ in *I*_*syn,*1_ given by (3). The phase-plane diagram is 2D. Because there is only one cell involved, we omit the subscript in the notation of the participating variables and parameters. The *v*- and *w*-nullclines are given, respectively, by

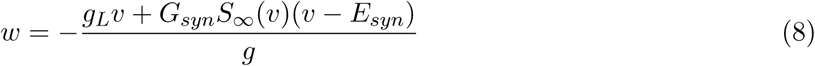

and

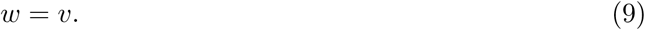

#### 2.2.3 Graded network of two 2D cells: hyper-nullclines, fixed-points and dynamic phase-plane analysis

These networks are given by system (1)-(4). The phase-space diagram is 4D. The *v*_1_- and *v*_2_-nullsurfaces (obtained by making the current-balance equation for the corresponding nodes equal to zero) depend on different variables (the *v*_1_-nullsurface depends on *w*_1_ and *v*_2_ and the *v*_2_- nullsurface depends on *v*_1_ and *w*_2_). The *w*_1_- and *w*_2_-nullsurfaces are planes given by *w*_1_ = *v*_1_ and *w*_2_ = *v*_2_, respectively. By substituting into the corresponding current-balance equations and rearranging terms we obtain the following equations describing curves in the *v*_1_-*v*_2_ plane

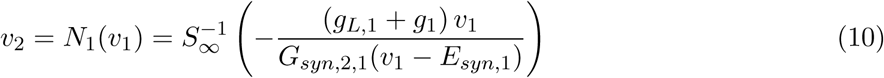

and

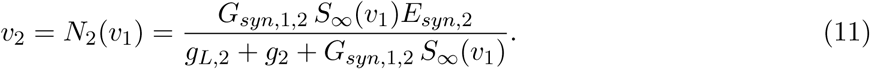

These are extensions of the nullclines (5) and (6) for the networks of 1D passive cells. Their inter-section 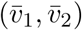 give the *v*_1_- and *v*_2_-coordinates of the 4D fixed-points 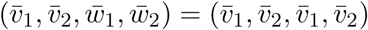. However, they are not nullclines, but projections of hyper-nullsurfaces onto the *v*_1_-*v*_2_ plane. We refer to them as hyper-nullclines. For the hybrid networks having one 2D and one 1D cell we set *g*_2_ = 0 in (11).

### 2.3 Bifurcation diagrams

As we mentioned in the previous section, the fixed-points are the intersections between the nullclines (for 2D systems) or the hyper-nullclines (for 3D and 4D systems). To determine the stability of the fixed-points we calculate the eigenvalues of the corresponding linearized system. For the 2D system of two 1D passive cells, the eigenvalues are easily calculated (see Appendix A). The expressions of the eigenvalues for the other considered networks (3D or 4D) are much more extensive and we will not show them in this work.

In all systems we can study the eigenvalue expressions when the parameter values vary, and we determine the existence of static bifurcations (such as, pitchfork and saddle-node) and dynamic bifurcations (for example, Hopf bifurcation) [77]. If a Hopf bifurcation exists, we calculate the first Lyapunov coefficient with the MATLAB package MatCont [78], to determine the direction and stability of the emerging branch of cycles.

Considering the bifurcations of the fixed-points, we construct bifurcation diagrams in several parameter spaces determining regions with different dynamical scenarios. In particular, we can determine parameter values in which stable limit cycles exist.

### 2.4 Conductance-based models

Primarily for illustrative purposes, in some of our simulations we used biophysical (conductance-based) models [79, 80] to describe the subthreshold dynamics of neurons having one resonant and one fast amplifying currents. The current balance equation is given by

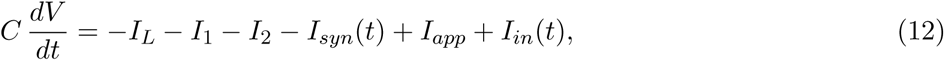

where *V* is the membrane potential (mV), *t* is time (ms), *C* is the membrane capacitance (*μ*F/cm^2^), *I*_*app*_ is the applied bias (DC) current (*μ*A/cm^2^), *I*_*in*_(*t*) is a time-dependent input current (*μ*A/cm^2^), *I*_*L*_ = *G*_*L*_ (*V* − *E*_*L*_) is the leak current, and *I*_*j*_ = *G*_*j*_ *x*_*j*_ (*V* − *E*_*j*_) are generic expressions for ionic currents (*j* = 1, 2) with maximal conductance *G*_*j*_ (mS/cm^2^) and reversal potentials *E*_*j*_ (mV) respectively. The gating variables obey kinetic equations of the form

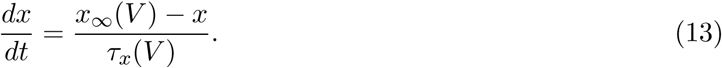

where *x*_*j,∞*_(*V*) and *τ*_*j,x*_(*V*) are the voltage-dependent activation/inactivation curves and time constants respectively. The former are given by

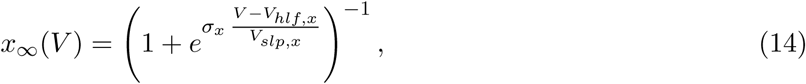

where *V*_*hlf,x*_ and *V*_*slp,x*_ > 0 are constants and the sign of *σ*_*x*_ indicates whether the curve describes an activation (*σ*_*x*_ < 0) or inactivation (*σ*_*x*_ > 0) process. In this paper we use voltage-independent time constants *τ*_*j,x*_. This assumption is mostly for simplicity since we are focusing on the subthreshold voltage regime where the time constants are typically slowly varying functions of *V*.

The ionic currents *I*_*j*_ we consider here are persistent sodium, *I*_*Nap*_ = *G*_*Nap*_ *p*_*∞*_(*V*) (*V* − *E*_*Na*_), hyperpolarization-activated, mixed-cation, inward (or h-), *I*_*h*_ = *G*_*h*_ *r* (*V* − *E*_*h*_) and slow-potassium (M-type) *I*_*Ks*_ = *G*_*Ks*_ *q* (*V* − *Ek*).

### 2.5 Numerical simulations

The numerical solutions were computed by using the modified Euler method (Runge-Kutta, order 2) [81] with a time step *Δt* = 0.1 ms in MATLAB (The Mathworks, Natick, MA). Smaller values of *Δt* have been used to check the accuracy of the results.

## 3 Results

### 3.1 Two-cell networks of passive cells do not produce limit cycle oscillations

Here we discuss some basic results regarding networks consisting of two interconnected cells with no cellular resonance and no self-connections. We use system (1) with *g*_*k*_ = 0 (*k* = 1, 2). The linearized conductances are restricted to be *g*_*L,*1_ > 0 and *g*_*L,*2_ > 0. Negative values of these linearized conductances are possible from the linearization procedure [2, 3]. However, the lack of both nonlinearities and intrinsic (resonant) negative feedback effects would cause the fixed-points (if they exist) to be unstable. It is conceivable though that network stabilize the instabilities of the individual cells under certain constraints.

The mathematical structure of these networks is similar to the two-dimensional firing rate models of Wilson-Cowan type [5, 68, 69] (with no self-connections) and some of the results for these models qualitatively apply here. For example, two-cell networks with the same connectivity sign (excitation or inhibition) and excitatory-inhibitory networks without recurrent (self-) excitation cannot produce sustained oscillations. We refer the reader to these references for further details.

The equations for the *v*_1_- and *v*_2_-nullclines are given by eqs. (5) and (6), respectively. Expressions for the eigenvalues for the fixed-points are presented in the Appendix A.

#### 3.1.1 Inhibitory networks of passive cells do not produce oscillations

The *v*_1_- and *v*_2_-nullclines are quasi-linear for low values of the maximal synaptic conductances *G*_*in,*1,2_ and *G*_*in,*2,1_, respectively (Figs. 3-a1, -b1 and -c1) and therefore the network has a single fixed-point, which is stable. Increasing values *G*_*in,*1,2_ and *G*_*in,*2,1_ cause the corresponding nullclines to develop nonlinearities and additional fixed-points may emerge and be destroyed. These fixed-points can be either stable nodes or saddles (see Appendix A).

**Figure 2:**
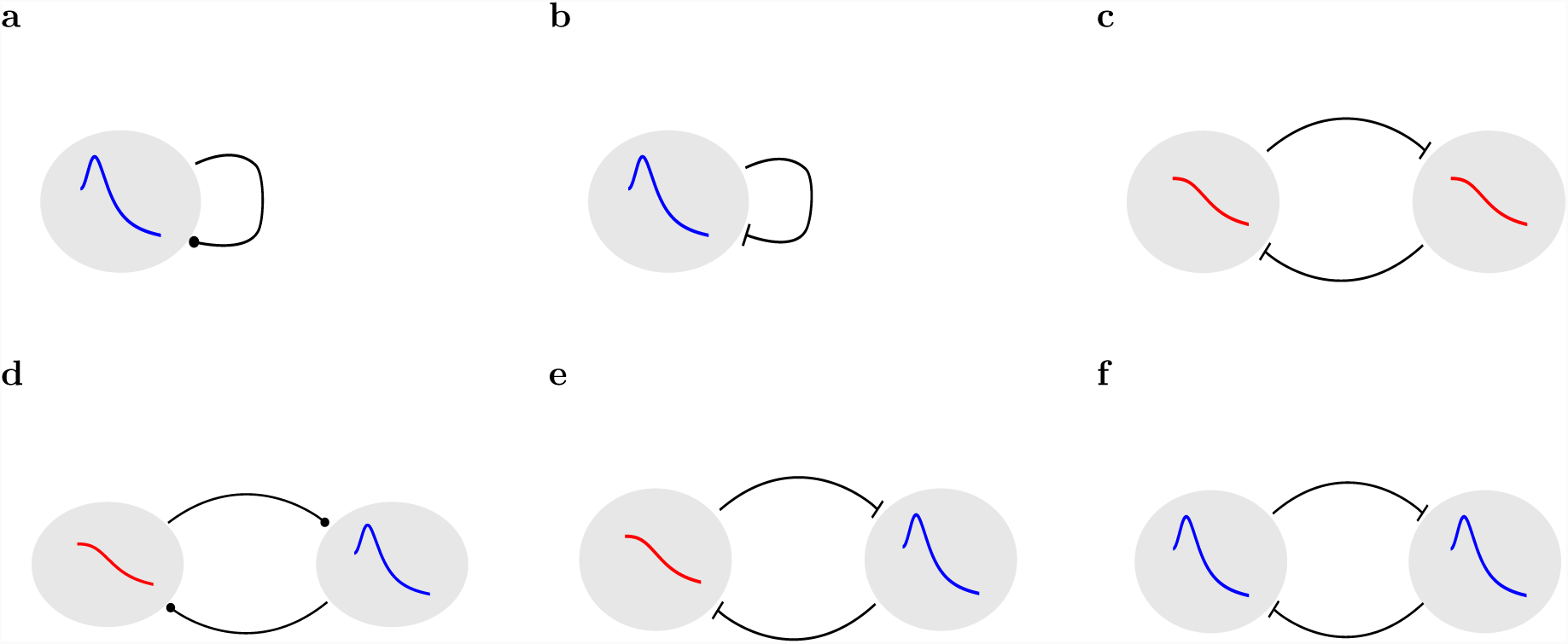
Network diagrams. **a.** Self-excited resonator (2D). **b.** Self-inhibited resonator (2D). **c.** Mutually inhibited passive cell network (1D/1D). **d.** Mutually inhibited resonator - passive cell network (2D/1D). **e.** Mutually excited resonator - passive cell network (2D/2D). e. Mutually inhibited resonator network (2D).

**Figure 3:**
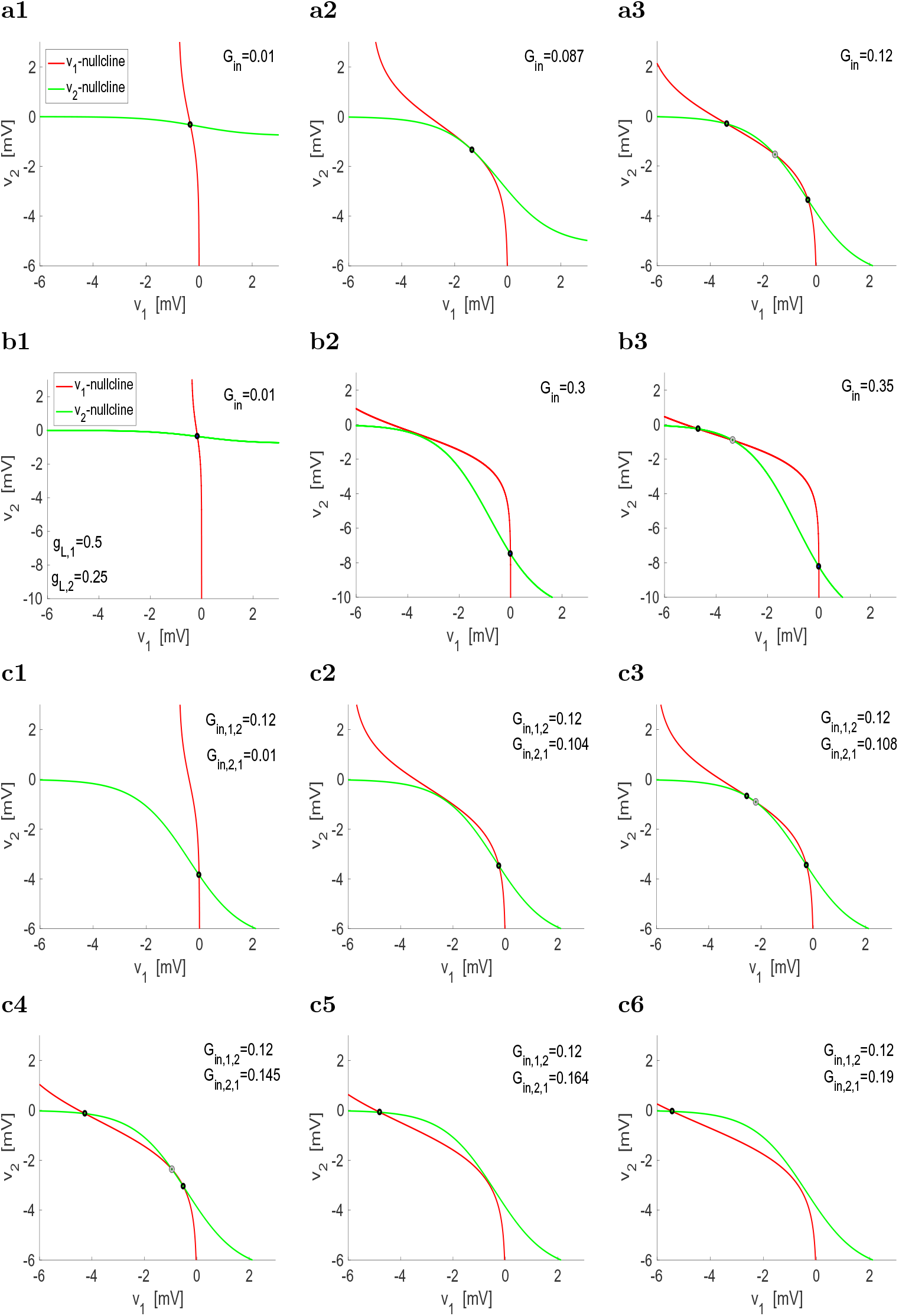
Reciprocally inhibitory networks of passive cells: phase-plane diagrams. The *v*_1_- and *v*_2_-nullclines are given by (5) and (6), respectively. Black dots indicate stable fixed-points (nodes) and gray dots indicate unstable fixed-points (saddles). **a** Identical cells and connectivity. The parameter *G*_*in*_ represents *G*_*in,*1,2_ = *G*_*in,*2,1_. **b.** Non-identical cells with identical connectivity. The parameter *G*_*in*_ represents *G*_*in,*1,2_ = *G*_*in,*2,1_. **c.** Identical cell with non-identical connectivity. We used the following parameter values: *g*_*L,*1_ = *g*_*L,*2_ = 0.25, *E*_*in,*_1 = *E*_*in,*_2 = −20, *v*_*hlf*_ = 0, *v*_*slp*_ = 1.

For homogeneous networks (*g*_*L,*1_ = *g*_*L,*2_, *G*_*in,*1,2_ = *G*_*in,*2,1_, *E*_*syn,*1_ = *E*_*syn,*2_ and identical functions *S*_*∞*_(*v*)), the stable fixed-point for low enough values of *G*_*in*_ (= *G*_*in,*1,2_ = *G*_*in,*2,1_) is symmetric (Fig. 3-a1). As *G*_*in*_ increases above some critical value (Fig. 3-a2), a pitchfork bifurcation occurs. The symmetric fixed-point becomes unstable and two (non-symmetric) stable fixed-points are created (Fig. 3-a3). As *G*_*in*_ increases further, the fixed-points move along the nullclines, but the system remains bistable (not shown).

For heterogeneous networks, bistability is created and terminated by saddle-node bifurcations (Figs. 3-b and -c). Fig. 3-c illustrates this for a network of identical cells with heterogeneous connectivity as the ratio *G*_*in,*1,2_*/G*_*in,*2,1_ decreases (*G*_*in,*2,1_ increases with *G*_*in,*1,2_ fixed). For low enough values of *G*_*in,*2,1_ cell 2 is inhibited (Figs. 3-c1 and c2). As *G*_*in,*2,1_ increases a stable and an unstable fixed-points are created (Fig. 3-c3). The stable fixed-point corresponds to an inhibited state for cell 1. The basin of attraction (measured as the distance between the stable and unstable fixed-points) is larger for the fixed-point corresponding to cell 2 being inhibited (stable fixed-point to the right). As *G*_*in,*2,1_ increases further, the unstable fixed-point moves closer to this fixed-point (Fig. 3-c4). The two of them eventually collide and disappear as *G*_*in,*2,1_ continues to increase leaving the inhibited cell 1 as the only steady state (Fig. 3-c5 and c6). Qualitatively similar dynamics are obtained for intrinsically heterogeneous networks with homogeneous/heterogeneous connectivity (not shown).

#### 3.1.2 Excitatory-inhibitory networks of passive cells produce damped oscillations for balanced levels of excitation and inhibition

We illustrate this in Fig. 4. From eq. (22) in the Appendix A, for fixed values of *G*_*ex,*1,2_ the eigenvalues are both real and negative for low enough values of *G*_*in,*2,1_ (Fig. 4-a). As *G*_*in,*2,1_ increases, the radicand decreases and eventually becomes negative leading to complex eigenvalues (Fig. 4-b). As *G*_*in,*2,1_ increases further, the effect on *F*_*v*_2 is balanced the effect on *F*_*v*_1, and the eigenvalues become real again (Fig. 4-c). These eigenvalues never loose stability and therefore, no limit cycle oscillations exist in these networks.

**Figure 4:**
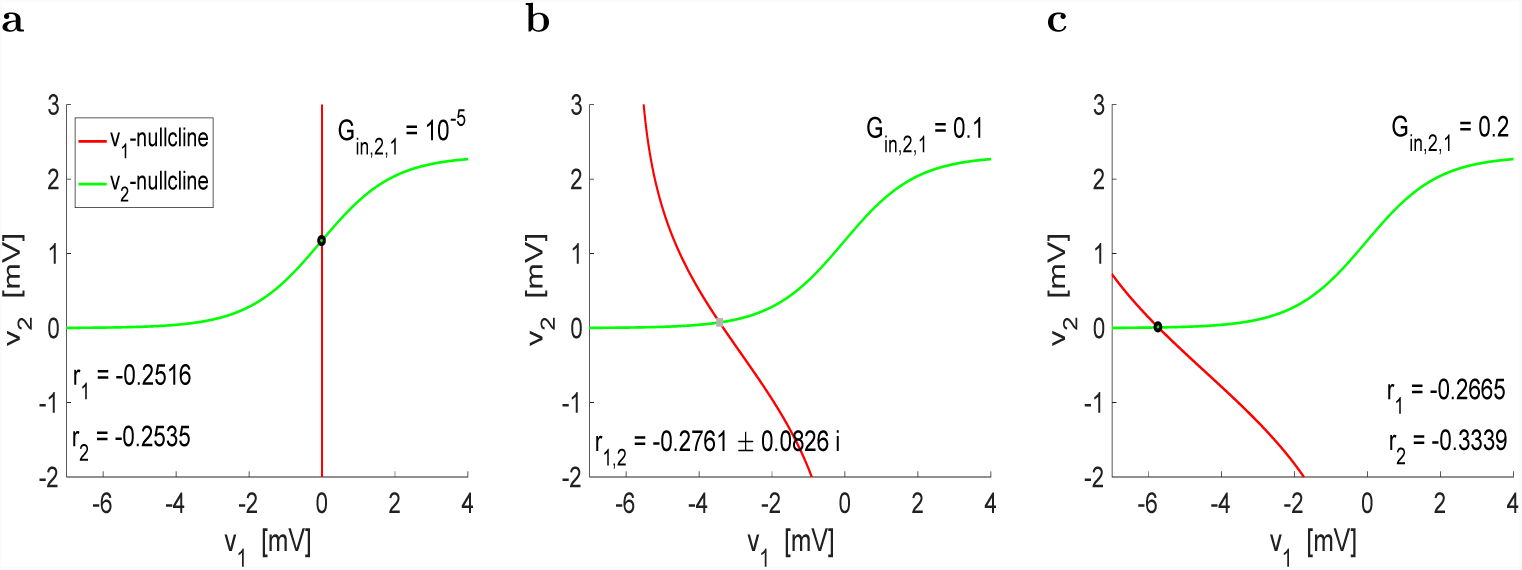
Excitatory-inhibitory networks of passive cells: phase-plane diagrams. The *v*_1_- and *v*_2_-nullclines are given by (5) and (6), respectively. Black dots indicate stable nodes and gray dots indicate stable foci. For fixed values of *G*_*ex,*1,2_ = 0.01, as *G*_*in,*2,1_ increases, the fixed-point transitions from stable nodes to stable foci and back to stable nodes. We used the following parameter values: *g*_*L,*1_ = *g*_*L,*2_ = 0.25, *E*_*in,*_1 = *E*_*ex,*_2 = 60, *v*_*hlf*_ = 0, *v*_*slp*_ = 1.

### 3.2 Self-excited resonators can produce limit cycle oscillations and their frequency monotonically depends on the resonator’s resonant frequency

The self-excited resonator model is given by system (1)-(2) where *v*_*k*_ = *v*_*j*_ in *I*_*syn,k*_ (3). Because there is only one cell involved, we omit the subscripts in the notation of the variables and parameters. The nullclines of the phase-plane diagrams are given by (8) and (9). The individual resonator does not oscillate.

Self-excitation is the simplest mechanism of network oscillation amplification of a resonator. Mathematically, a self-excited resonator has the same structure as individual resonator+amplifying current models (e.g., *I*_*h*_+ *I*_*Nap*_ or *I*_*Ks*_+*I*_*Nap*_), which are able to produce sustained oscillations for large enough amplification levels (Fig. 1) [9]. In both types of models the activation of the amplifying component (*I*_*syn*_ and *I*_*Nap*_) is instantaneous (or very fast), the shapes of their activation curves are similar, and the reversal potentials (*E*_*Na*_ and *E*_*ex*_) are above the resting potential. Models having *I*_*h*_ or *I*_*Ks*_ as the only active ionic currents are quasi-linear resonators [9]. Therefore, it is not surprising that self-excited linear resonators are able to produce oscillations given that resonant+amplifying models can do so. However, since resonance and amplification belong to different levels of organization in self-excited resonators, we can dissociate these two effects and investigate the effects of the resonant frequency of the individual neurons on the oscillation frequency, which we cannot do in individual cells.

#### 3.2.1 Self-excited resonators can produce sustained (limit cycle) oscillations for appropriate balances among the resonance, amplification and excitation levels

Geometrically, increasing values of the excitatory maximal conductance *G*_*ex*_ create nonlinearities of cubic type in the phase-planed diagram (Fig. 5). In single neurons this type of nonlinearities are typically created by amplifying gating variables (e.g., *I*_*Nap*_) in the presence of resonant gating variables (e.g., *I*_*h*_ or *I*_*Ks*_) [9, 11] (see also [5, 7]).

**Figure 5:**
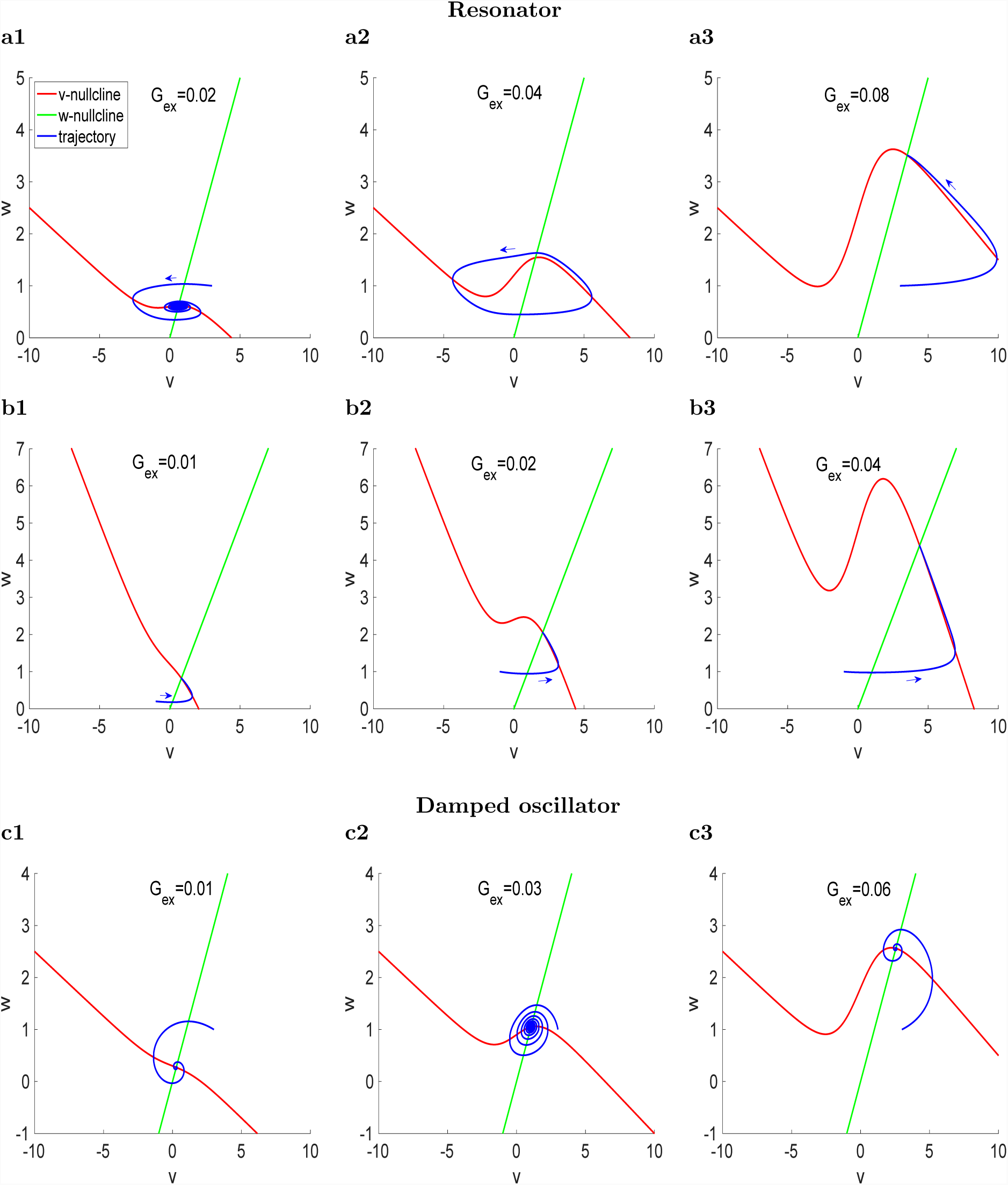
Self-excited (linear) resonators can produce sustained (limit cycle) oscillations, while self-excited damped-oscillators may fail to produce sustained oscillations. Phase-plane diagrams for representative parameter values. The *v*- and *w*-nullclines are given by (8) and (9), respectively. The fixed-point for the uncoupled (linear) system is a stable node (*f*_*nat*_ = 0) in panels a and b and a stable focus in panels c (*f*_*nat*_ ∼ 48.9). **a.** *f*_*res*_ ∼ 17.6 for *g*_*L*_ = 0.25, *g* = 1 and *τ* = 100. **b.** *f*_*res*_ ∼ 10.4 for *g*_*L*_ = 0.25, *g* = 0.25 and *τ* = 100. **c.** *f*_*res*_ ∼ 55.2 for *g*_*L*_ = 0.25, *g* = 1 and *τ* = 10. We used the following additional parameter values:, *E*_*ex*_ = 60, *v*_*hlf*_ = 0, *v*_*slp*_ = 1.

Fig. 5-a illustrates the effects of increasing values of *G*_*ex*_ when the linearized resonant conductance *g* (= 1) is much larger than the linearized leak conductance *g*_*L*_ (= 0.25) for a resonator (with no intrinsic damped oscillations when *G*_*ex*_ = 0). For low values of *G*_*ex*_, the coupled cell shows damped oscillations as the cubic-like nonlinearities of the *v*-nullcline begin to develop (panel a1). Limit cycle oscillations emerge as *G*_*ex*_ increases further (panel a2) and disappear when the fixed-point moves to the right branch of the cubic-like *v*-nullcline for larger values of *G*_*ex*_ and regains stability (panel a3). As *G*_*ex*_ increases within the oscillatory range the amplitude increases and a time scale separation between the participating becomes more prominent, generating, for large enough values of *G*_*ex*_, oscillations of relaxation type (Fig. 6-a).

**Figure 6:**
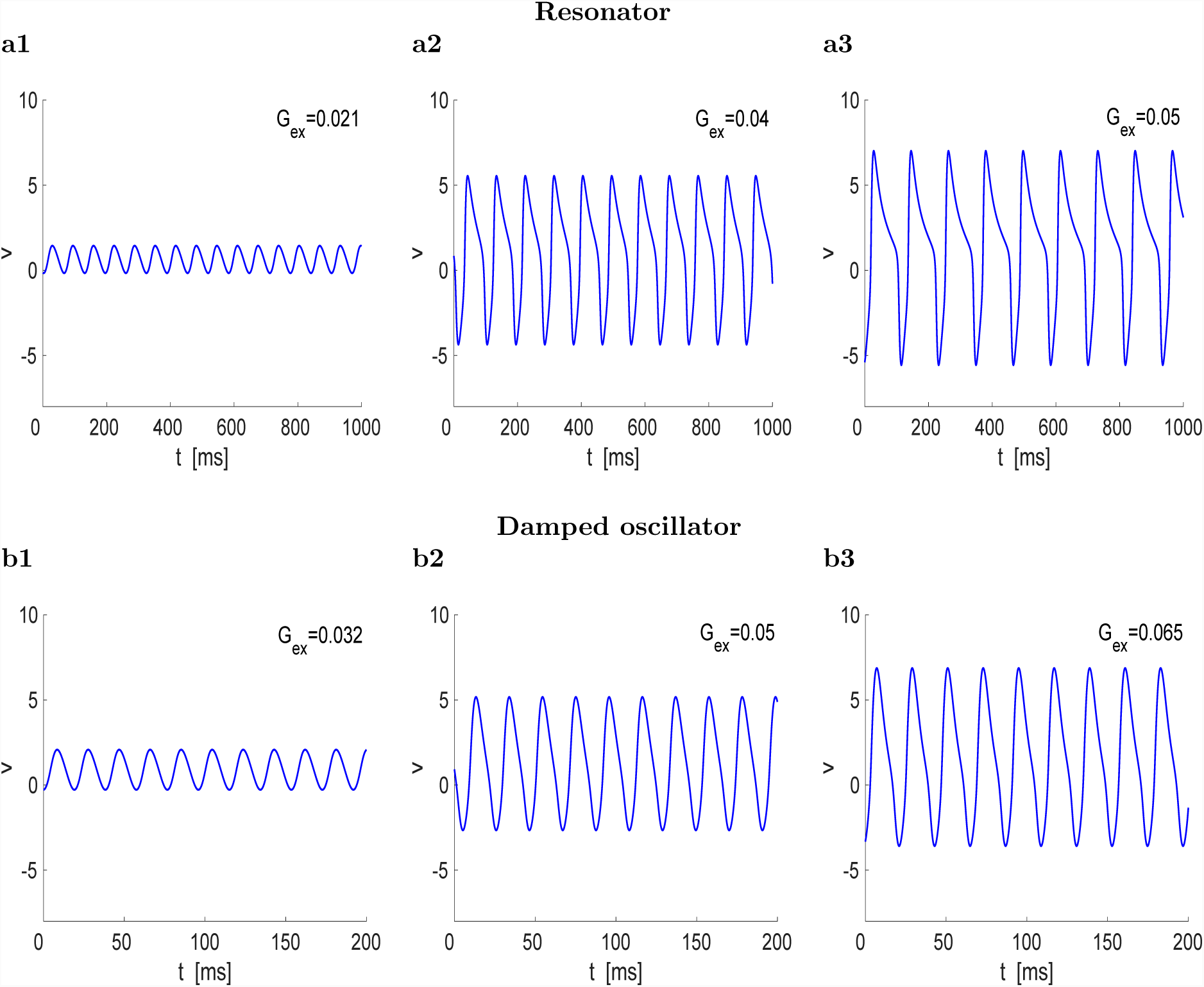
Development of relaxation oscillations in self-excited resonators, but not necessarily in self-excited damped-oscillators. (a) Self-excited resonator. Parameter values are as in Fig. 5-a: *f*_*res*_ ∼ 17.6 and *f*_*nat*_ = 0 for *g*_*L*_ = 0.25, *g* = 1 and *τ* = 100. Oscillations are generated/terminated by Hopf bifurcations for low (supercritical) and high (subcritical) values of *G*_*ex*_. (a1) *f*_*ntw*_ = 15.5 for *G*_*ex*_ = 0.021. (a2) *f*_*ntw*_ = 11.1 for *G*_*ex*_ = 0.04. (a3) *f*_*ntw*_ = 8.5 for *G*_*ex*_ = 0.05. (b) Self-excited damped-oscillator. Parameter values are as in Fig. 5-c, except for *g* that is larger to produce sustained oscillations: *f*_*res*_ ∼ 59.8 and *f*_*nat*_ = 53.8 for *g*_*L*_ = 0.25, *g* = 1.2 and *τ* = 10. (b1) *f*_*ntw*_ = 52.1 for *G*_*ex*_ = 0.032. (b2) *f*_*ntw*_ = 48.5 for *G*_*ex*_ = 0.05. (b3) *f*_*ntw*_ = 45.5 for *G*_*ex*_ = 0.065. We used the following additional parameter values: *E*_*ex*_ = 60, *v*_*hlf*_ = 0, *v*_*slp*_ = 1.

Fig. 5-b illustrates that oscillations are not generated when the *g* (= 0.25) to *g*_*L*_ (= 0.25) ratios are relatively low. The cubic-like nonlinearities are still developed for high enough values of *G*_*ex*_ (panels b2 and b3), but the fixed-point is located on the right branch of the *v*-nullcline where the fixed-point is stable, and moves further away from the knee as *G*_*ex*_ increases. The amplification still happens, but it leads directly to depolarization block without oscillations. Similar behavior was observed when the fixed-point of the isolated cell is a stable focus instead of a stable node. However, oscillations can be restored by increasing the value of *v*_*hlf*_, which moves the fixed-point to the middle branch where it loses stability (not shown).

The transition of a resonator to a damped oscillator can be achieved by decreasing the value of *τ* [3]. Contrary to intuition, the presence of damped oscillations in the cell does not necessarily generate sustained oscillations in the self-excited network (Fig. 5-c). When it happens, the time scale separation is smaller than for the resonator and therefore relaxation oscillations are more difficult to obtain (Fig. 6-b).

#### 3.2.2 The intrinsic resonant frequency controls the network oscillations frequency

Self-excited resonators are the simplest models where we can investigate the effects that changes on the resonant frequency (*f*_*res*_) of the individual non-oscillatory cells have on the network oscillation frequency (*f*_*ntw*_). The resonator parameters that control *f*_*res*_ (26) also control the values of other attributes of the impedance profile *Z*(*f*) such as the maximal impedance *Z*_*max*_ (27). In order to establish the effects of *f*_*res*_ on *f*_*ntw*_ it is necessary change the model parameters in such a way as to cause the minimal possible changes on the shape of *Z*(*f*) [41]. In the ideal situation, changes in *f*_*res*_ would be accompanied only by a translation of *Z*(*f*). This is not possible for 2D linear models, but it is possible to change the model parameters in a balanced way so that *f*_*res*_ changes, but *Z*_*max*_ remains constant [41]. In this way the impedance profiles are displaced with minimal changes in their shape (Fig. 7-a).

**Figure 7:**
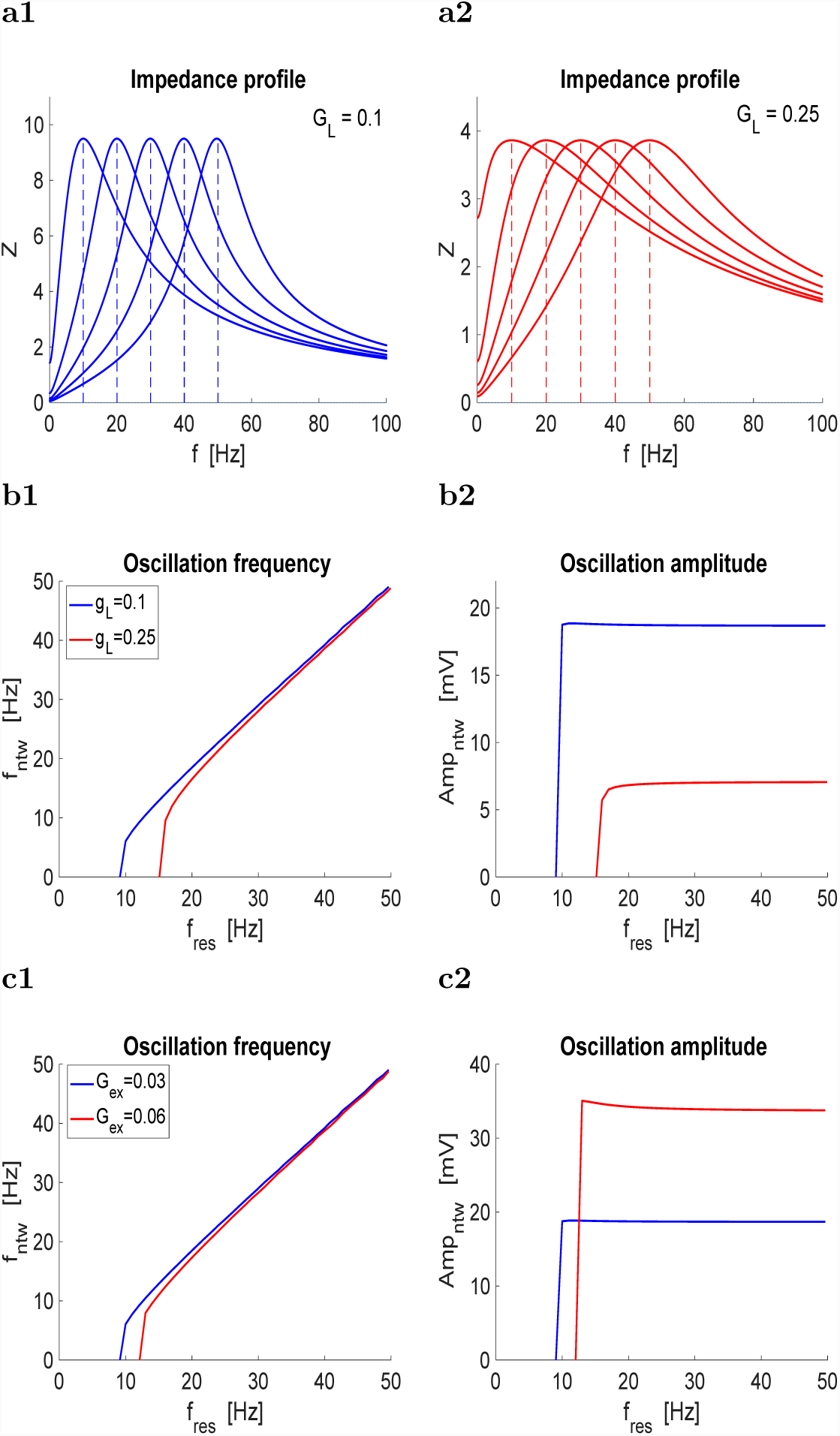
Oscillations in self-excited resonators: the intrinsic resonant frequency controls the network frequency. a. Representative resonator impedance profiles with different resonant frequencies (*f*_*res*_) and the same maximal impedance: *Z*_*max*_ ∼ 9.5 (a1) and *Z*_*max*_ = 3.9 (a2). **b.** Network oscillation frequency (b1) and amplitude (b2) as a function of *f*_*res*_ for representative values of *g*_*L*_. **c.** Network oscillation frequency (c1) and amplitude (c2) as a function of *f*_*res*_ for representative values of *G*_*ex*_. We used the following parameter values: *E*_*ex*_ = 60, *v*_*hlf*_ = 0, *v*_*slp*_ = 1.

Figs. 7-b and -c show that increasing values of *f*_*res*_ directly affect *f*_*ntw*_ (Figs. 7-b1 and -c1) with minimal changes in the oscillation amplitude (Figs. 7-b2 and -c2). The onset of oscillations occurs for lower values of *f*_*res*_ the lower *g*_*L*_ (Figs. 7-b1) and *G*_*ex*_ (Figs. 7-c1). As expected, the oscillations are more amplified the lower *g*_*L*_ (Figs. 7-b2) [3, 9] and the higher *G*_*ex*_ (Figs. 7-c2).

#### 3.2.3 Self-inhibited resonators do not produce sustained (limit cycle) oscillations

The primary effect of self-inhibition is to attenuate signals. By this we mean to reduce the equilibrium values (Fig. 8-a), to reduce the amplitude of the damped oscillations (measured, for example, as the difference between a maximum and the consecutive minimum), or to transform damped oscillations into trajectories of a stable node (Fig. 8-b).

**Figure 8:**
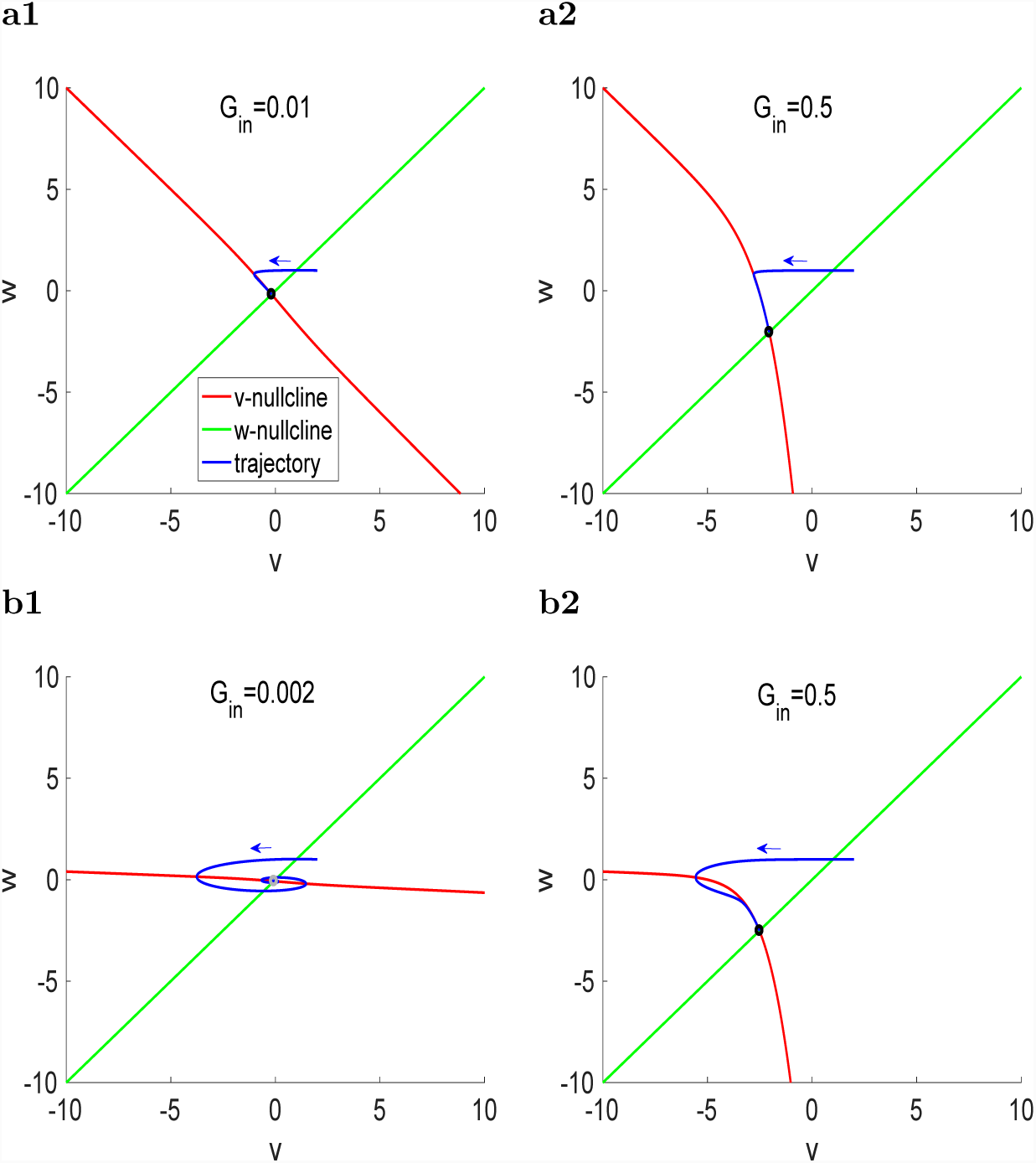
Self-inhibited resonant cells: phase-plane diagrams for representative parameter values. The *v*- and *w*-nullclines are given by (8) and (9), respectively. Black dots indicate stable nodes and gray dots indicate stable foci. **a.** *g*_*L*_ = 0.25. The fixed-point for the uncoupled system is a stable node. **b.** *g*_*L*_ = 0.01. The fixed-point for the uncoupled system is a stable focus. We used the following parameter values: *g* = 0.25, *τ* = 100, *E*_*in*_ = −20, *v*_*hlf*_ = 0, *v*_*slp*_ = 1.

The presence of self-inhibition generates nonlinearities in the *v*-nullcline, while the *w*-nullcline remains linear (as for the uncoupled system) (Figs. 8-a2 and -b2). The dependence of these nonlinear effects with the connectivity parameters is illustrated in Figs. 8. For low values of *G*_*in*_ the system is quasi-linear (Figs. 8-a1 and -b1) and the dynamics qualitatively remains as for the uncoupled system (not shown). For larger values of *G*_*in*_ the *v*-nullcline bends creating two quasi-linear portions (Figs. 8-a2 and -b2) with different slopes. For the connectivity parameters used in Figs. 8-a2 and -b2 (*G*_*in*_, *v*_*hlf*_ and *v*_*slp*_) the fixed-point is located on the branch of the *v*-nullcline with the steepest slope, which in some cases changes the properties of the fixed-point (Fig. 8-b). For higher values of *v*_*hlf*_ where the synaptic function moves to the right, effectively decreasing the connectivity, the *v*-nullcline remains nonlinear, but the fixed-point moves to the other portion of the *v*-nullcline (not shown).

#### 3.2.4 Self-excited low-pass filter cells do not produce sustained (limit cycle) oscillations

Resonance can be lost by various mechanisms [3]. One of them is having low enough values of the resonant conductance *g* (in the limit of *g* = 0 the coupled cell is 1D and therefore oscillations are not possible). Another one is having low-enough values of the time constant *τ*. Fig. 9-a illustrates how oscillations are lost as *τ* decreases. Note that the location of the fixed-point is independent of *τ*. Figs. 9-b and -c illustrate that oscillations cannot be recovered by decreasing *G*_*ex*_ (panel b) or increasing *v*_*hlf*_ (panel c) for the same value of *τ* as in panel a3. In both cases, these changes move the fixed-point to the middle branch of the *v*-nullcline, but it remains stable.

**Figure 9:**
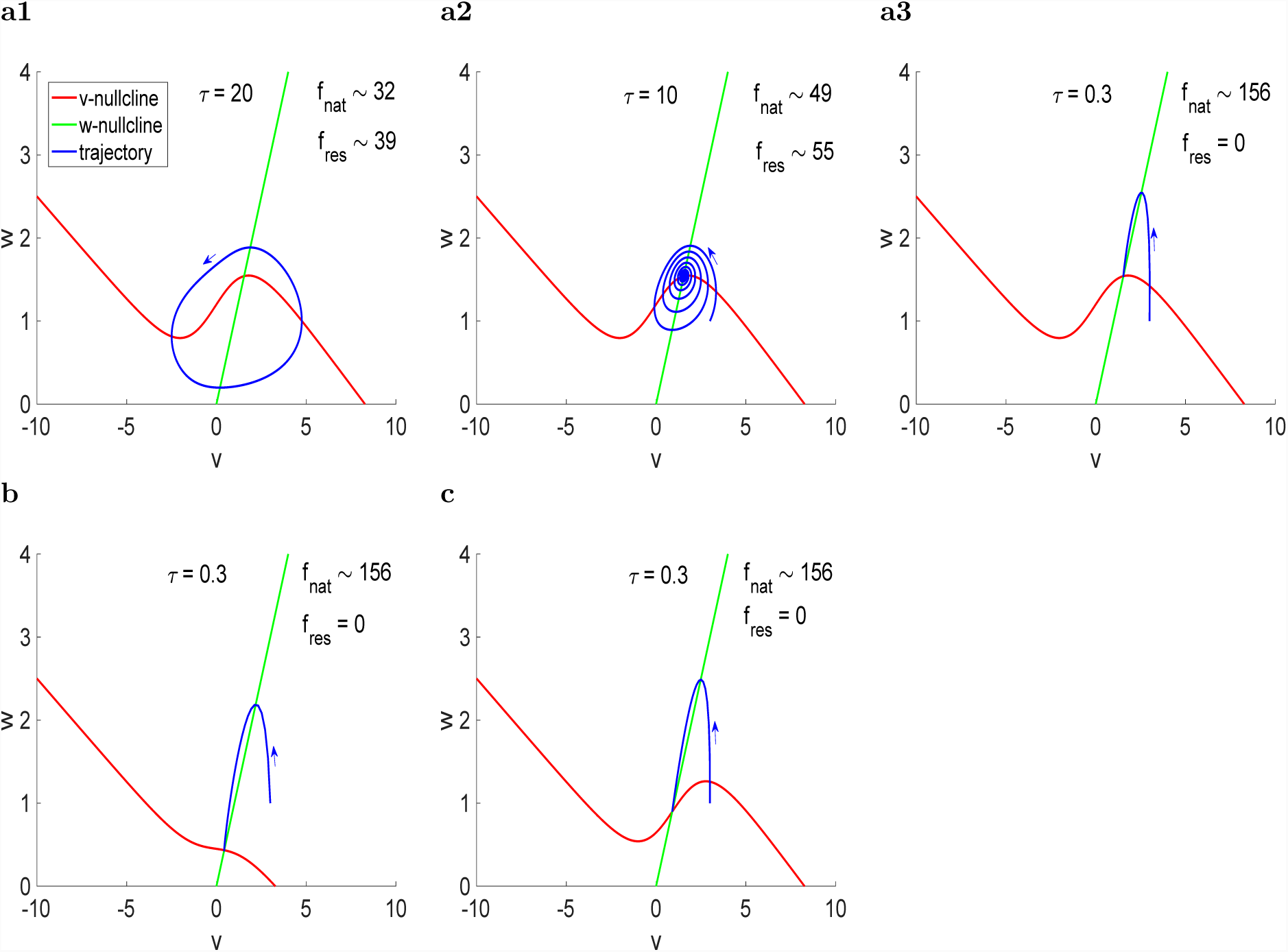
Self-excited 2D cells: phase-plane diagrams for representative parameter values. The *v*- and *w*-nullclines are given by (8) and (9), respectively. The quantities *f*_*nat*_ and *f*_*res*_ refer to the natural and resonant frequencies of the uncoupled cells. The fixed-point for the uncoupled system is a stable focus. **a.** *G*_*ex*_ = 0.04 and *v*_*hlf*_ = 0. **b.** *G*_*ex*_ = 0.015 and *v*_*hlf*_ = 0. **c.** *G*_*ex*_ = 0.04 and *v*_*hlf*_ = 1. We used the following parameter values: *g*_*L*_ = 0.25, *g* = 1, *E*_*ex*_ = 60, *v*_*slp*_ = 1.

### 3.3 Mutually inhibitory 2D/1D hybrid networks can generate sustained (limit cycle) oscillations and their frequency monotonically depends on the intrinsic resonant frequency

The hybrid networks we consider here consist of a linear resonator (2D, cell 1) and a passive cell (1D, cell 2) reciprocally inhibited through graded synapses. We use system (1)-(2) with *g*_1_ > 0 and *g*_2_ = 0 and the additional description of the synaptic connectivity presented in Section 2.

These networks can be thought of as two “overlapping” circuits, non of each able to produce oscillations on their own: the linear 2D resonator used in Section 3.2 and the reciprocally inhibited passive cells discussed in Section 3.1. The oscillations result from the combined activity of these two “sub-circuits” where the mutually-inhibitory component acts as an amplifier of the resonant component.

For our analysis we represent the dynamics of these 3D networks using projections of the 3D phase-space (for *v*_1_, *w*_1_ and *v*_2_) onto the *v*_1_ - *v*_2_ plane and use the hyper-nullclines (10)-(11) (*g*_2_ = 0) defined in Section 2.2 (e.g., Fig. 10, left columns). In order to relate the dynamics of the hybrid networks to these of the mutually inhibitory passive cells (Section 3.1) we include in the phase-plane diagrams the *v*-nullcline for cell 1 (dashed-red curve) for *g*_1_ = 0 (no resonant gating variable).

**Figure 10:**
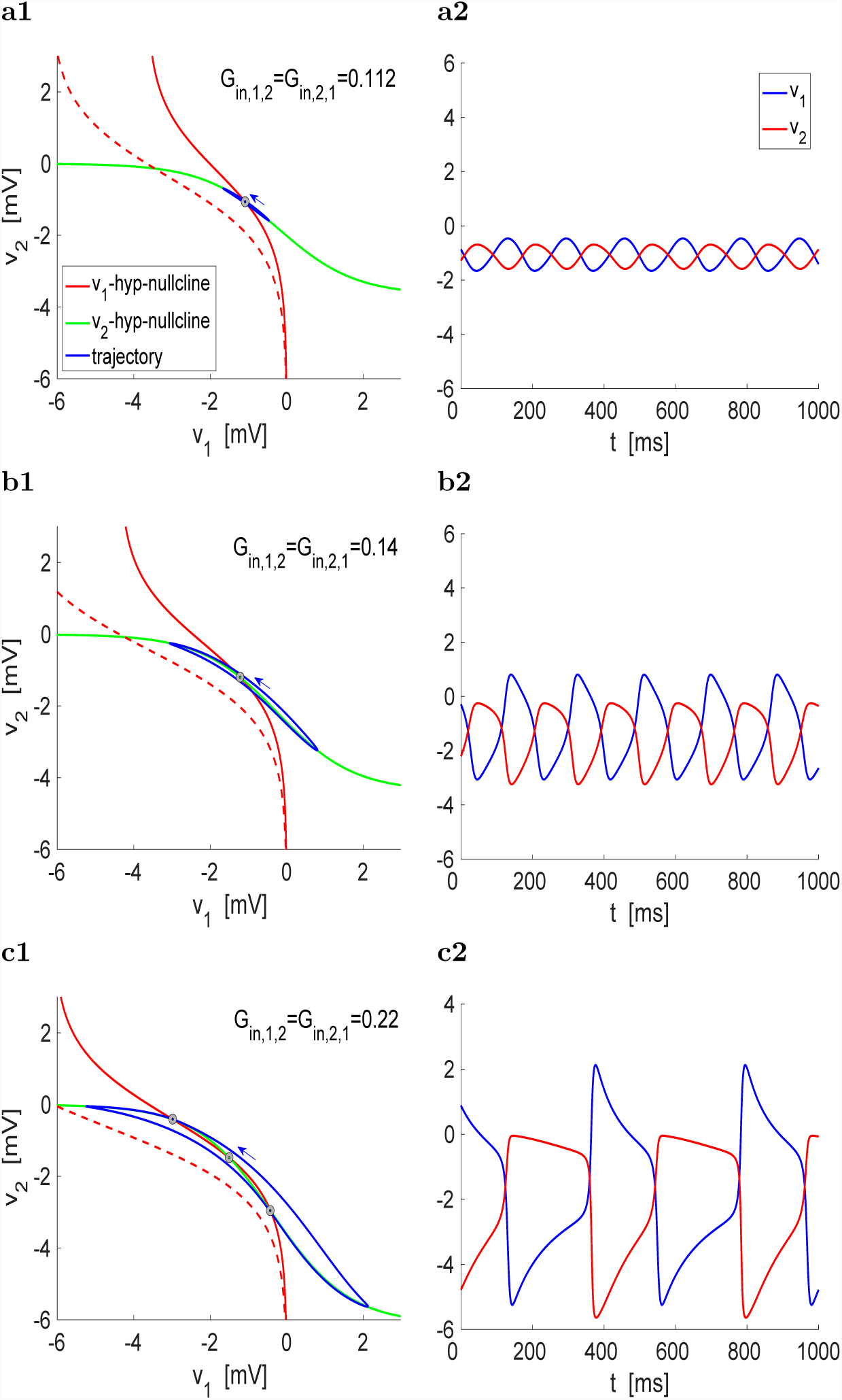
Oscillations generated in mutually inhibited hybrid 2D-1D networks. Cell 1 is a resonator with *f*_*res*_ = 10.4 (*f*_*nat*_ = 0) and cell 2 is a passive cell. **Left.** Phase-plane diagrams. The *v*_1_- and *v*_2_-hyper-nullclines are given by (10) and (11), respectively. Black dots indicate stable nodes and gray dots indicate unstable foci. The dashed red curve represents the *v*_1_ nullcline for cell 1 for *g*_1_ = 0 (no resonant gating variable). **Right.** Voltage traces (curves of *v*_1_ and *v*_2_ as a function of *t*). (a) *G*_*in,*1,2_ = *G*_*in,*2,1_ = 0.112. The network frequency is *f*_*ntw*_ = 6.1. (b) *G*_*in,*1,2_ = *G*_*in,*2,1_ = 0.14. The network frequency is *f*_*ntw*_ = 5.4. (c) *G*_*in,*1,2_ = *G*_*in,*2,1_ = 0.22. The network frequency is *f*_*ntw*_ = 2.4. We used the following parameter values: *g*_1_ = 0.25, *g*_*L,*1_ = 0.25, *g*_*L,*2_ = 0.5, *τ* = 100, *E*_*in*_ = −20, *v*_*hlf*_ = 0, *v*_*slp*_ = 1.

#### 3.3.1 Oscillations can be generated in 2D/1D hybrid networks and are amplified with increasing levels of mutual inhibition

Fig. 10 shows the oscillations generated in these networks for values of *G*_*in*_ (= *G*_*in,*1,2_ = *G*_*in,*2,1_) that increase from top to bottom. Because the networks are mutually inhibited, these oscillations are not synchronized in-phase. They are created in a supercritical Hopf bifurcation (Fig. 13-a1) and therefore they have small amplitude and are sinusoidal-like for small enough values of *G*_*in*_ (Fig. 10-a). The effect of the resonant gating variable *w*_1_ is to bring the fixed-point of the mutually inhibitory (non-oscillatory) 1D/1D system (intersection between the dashed-red and green curves) to the oscillatory region where the *v*_2_ hyper-nullcline (green curve) is non-linear.

The oscillations amplitude increases with increasing values of *G*_*in*_ as the limit cycle trajectories evolve in small vicinities of the *v*_2_ hyper-nullcline (Fig. 10-b and -c). This amplification is accompanied by the development of a separation of time scales. For large enough values of *G*_*in*_ the oscillations are of relaxation-type (Fig. 10-c). This partially reflects the time constant of the resonators, which needs to be slow enough, but it is a network effect since linear models do not display sustained oscillations.

For low values of *G*_*in*_ within the oscillatory regime, the network has only one fixed-point. As *G*_*in*_ increases, additional fixed-points are created (Figs. 10-c1 and 13-c1) in a Pitchfork bifurcation (Fig. 13-a1), but they are not stable and they do not obstruct the presence of oscillations. However, as *G*_*in*_ increase further, these new fixed-points become stable by subcritical Hopf bifurcations and coexist as attractors with the limit cycle (Fig. 13-a1). The oscillations are abruptly terminated when the stable limit cycle collides with an unstable limit cycle generated in one of the mentioned subcritical Hopf bifurcations (Fig 13-c1). Without oscillations the attractors that remain in the network are the fixed-points corresponding to one of the cells inhibited (Fig. 13-a1 and c1).

Similarly to the mutually inhibited passive cells discussed above, for other parameter regimes the pitchfork bifurcation can be transformed into a saddle-node bifurcation (Fig. 13-a2) without causing significant qualitative changes to the network dynamics (Fig. 13-c2).

Fig. 13-b illustrates that existence of network oscillations requires balanced combinations of *G*_*in*_ and *g*_1_.

The generation of oscillations requires certain heterogeneity in the underlying mutually inhibitory 1D/1D system. For the oscillations in Fig. 10 *g*_*L,*1_ = 0.25, *g*_*L,*2_ = 0.5. This has been observed also for the related system studied in [67]. Oscillations are not possible for the hybrid 2D/1D network when *g*_*L,*1_ = *g*_*L,*2_ and *C*_1_ = *C*_2_ unless there is heterogeneity in the synaptic connectivity (*G*_*in,*2,1_ > *G*_*in,*1,2_) (not shown).

#### 3.3.2 Development of relaxation oscillations for large mutual inhibition levels

In order to understand the mechanisms of generation of network oscillations and their properties in terms of the model parameters it is useful to consider the *v*_1_-nullsurface

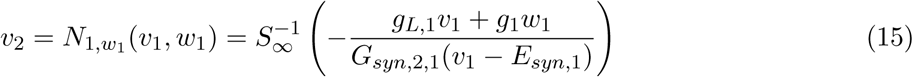

parametrized by constant values of *w*_1_, *N*_1*c*_(*v*_1_) = *N*_1*c*_(*v*_1_, *c*), and track the motion of the trajectory as time progresses and the values of *w*_1_ change. This will cause the *v*_1_-hyper-nullclines in Fig. 10 to move as the trajectory evolves following the dynamics of *w*_1_. For the second cell the curve (12) for *g*_2_ = 0 is time-independent and therefore remains fixed. Note that eq. (15) is eq. (10) before *w*_1_ is substituted by *v*_1_.

In order to uncover the presence of nonlinearities of cubic type and to further capture the effect of the model’s geometric properties that give rise to the different types of oscillations we use an adapted version (Fig. 11) of the phase-plane diagram discussed in Fig. 10 where the hyper-nullclines and trajectories are referred to the *v*_2_-hyper-nullcline. In the adapted phase-plane diagram, the *v*_2_ hyper-nullcline (green curve) is the zero-level line and the *v*_1_-hyper-nullclines (red curves) are cubic-like. The dashed-red curves represent the maximum (lower curve) and minimum (upper curve) levels of *w*_1_ during the oscillations. The red curve moves in between these two dashed-red curves as the oscillation progresses. The intersections between the green line and the moving red curve generated “transient fixed-points”, which are not fixed-points of the 3D system, but they serve as targets for the evolution of the trajectories. Specifically, in the *v*_1_-*v*_2_ plane presented in Fig. 11 (left panels), trajectories move towards the transient fixed-points with negative slope and their speed depends on the distance between the moving red curve and the green line. The existence of oscillations imply that the local extrema of the dashed-red curves do not intersect the zero-level reference green line. We emphasize that this is not a standard phase-plane diagram and it captures only specific aspects of the dynamics.

**Figure 11:**
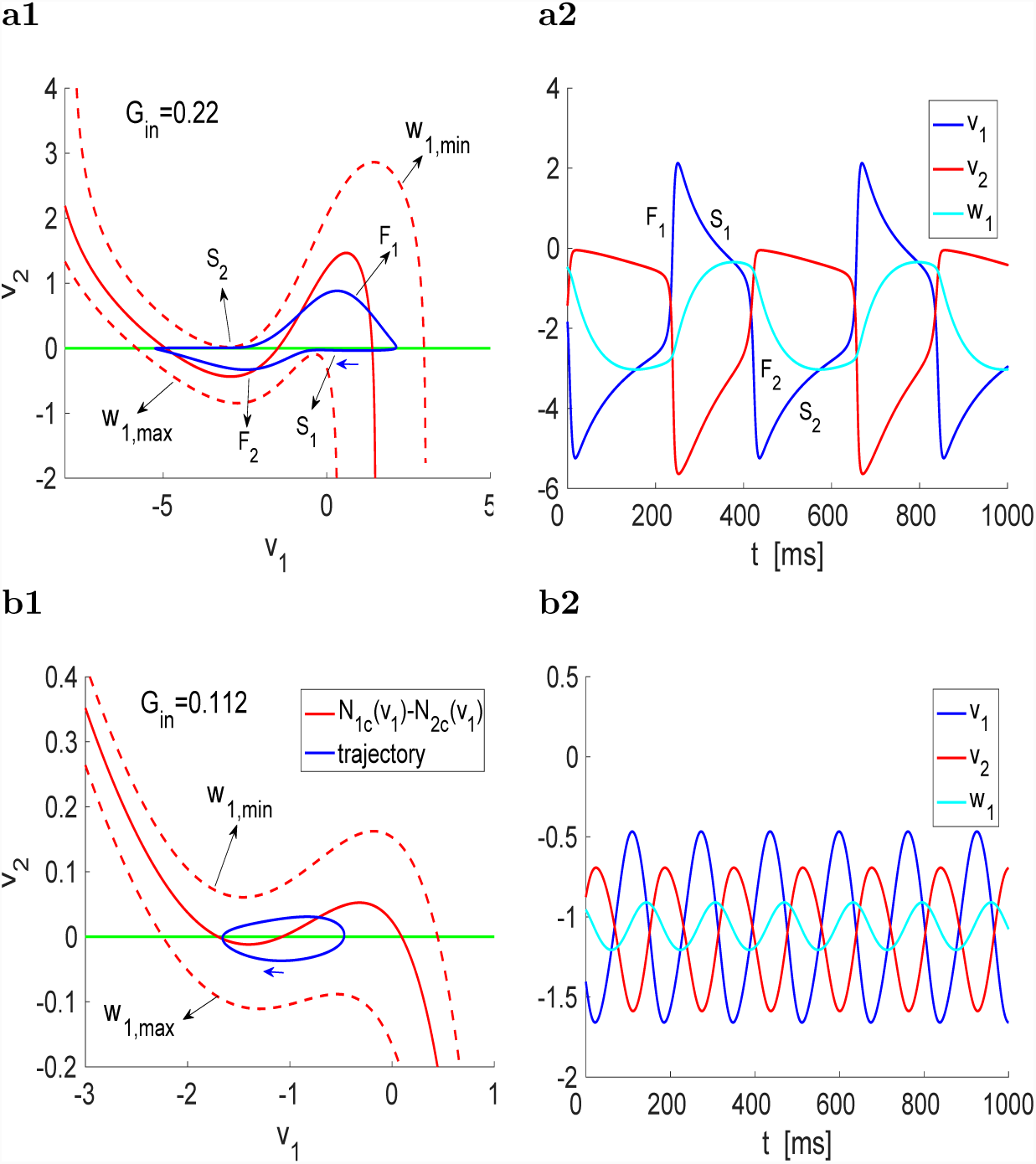
Development of relaxation oscillations for large mutual inhibition levels in hybrid 2D-1D networks. The parameter values correspond to Fig. 10. Cell 1 is a resonator with *f*_*res*_ = 10.4 (*f*_*nat*_ = 0) and cell 2 is a passive cell. The values of *G*_*in,*1,2_ = *G*_*in,*2,1_ are represented by *G*_*in*_. **Left.** Adapted phase-plane diagrams relative to the *v*_2_-hyper-nullcline *N*2_*c*_(*v*_1_) (green curve in the phase-plane diagrams in Fig. 10, left panels). The red lines are the differences between the *v*_1_- and *v*_2_-hyper-nullclines in Fig. 10 (left panels) parametrized by constant values of *w*_1_. The solid-red curve corresponds to the fixed-point 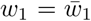. The dashed-red curves correspond to the maximal *w*_1,*max*_ (lower) and minimal *w*_1,*min*_ (upper) values of *w*_1_. The trajectories (blue curves) are also referred to the *v*_2_-hyper-nullcline *N*2_*c*_(*v*_1_). **Right.** Voltage traces for *v*_1_, *v*_2_ and *w*_1_. (a) Relaxation oscillations for *G*_*in,*1,2_ = *G*_*in,*2,1_ = 0.22. The network frequency is *f*_*ntw*_ = 2.4. (b) Oscillations with a uniform time scale for *G*_*in,*1,2_ = *G*_*in,*2,1_ = 0.112. The network frequency is *f*_*ntw*_ = 6.1.

Similarly the self-excited resonator discussed above, increasing amplification levels are characterized by more pronounced cubic-like nonlinearities. Here the amplification levels are provided by the levels of mutual inhibition that are measured in terms of the values of *G*_*in*_ (compare panels a1 and b1 in Fig. 11).

When *w*_1_ = *w*_1,*min*_ the red curve is at its highest level and the trajectory moves to the right (*F*_1_), towards the only transient fixed-point with relatively high speed (jump up). As this happens *w*_1_ increases, causing the red curve to shift down with the consequent motion of the transient fixed-point to the left. The variable *v*_1_ reaches its maximum when the trajectory crosses the transient fixed point and is forced to reverse direction (*S*_1_). As the red curve continues to shifts down, the stable and unstable transient fixed-points collide and disappear leaving only one transient fixed-point (on the leftmost side, for lower values of *v*_1_), which becomes the new target for the trajectory. The trajectory moves towards this target fixed-point, but it does so on a very slow time scale (*S*_1_) due to the ghost effect of the “defunct” fixed-points until it reaches the (jump down) region of fast motion (*F*_2_). The process repeats to complete the cycle through.

Relaxation oscillations are created when difference between the two local extrema on each dashed-red curve is large enough (well separated). This occus in Fig. 11-a, but not in Fig. 11-b, where the oscillations do not show any separation of time scales.

#### 3.3.3 The resonator’s intrinsic resonant frequency controls the network oscillations frequency

Similarly to the self-excited resonator networks discussed above, the functional role of cellular resonance is to determine the frequency of the network oscillations. This is illustrated in Fig. 12 for various representative parameter set values. We followed the same protocol as in Section 3.2.2 (Fig. 7): for each value of *f*_*res*_, the values of *g*_1_ and *τ*_1_ are balanced so to maintain *Z*_*max*_ constant. In all cases, *f*_*ntw*_ increases with increasing values of *f*_*res*_ (left panels).

**Figure 12:**
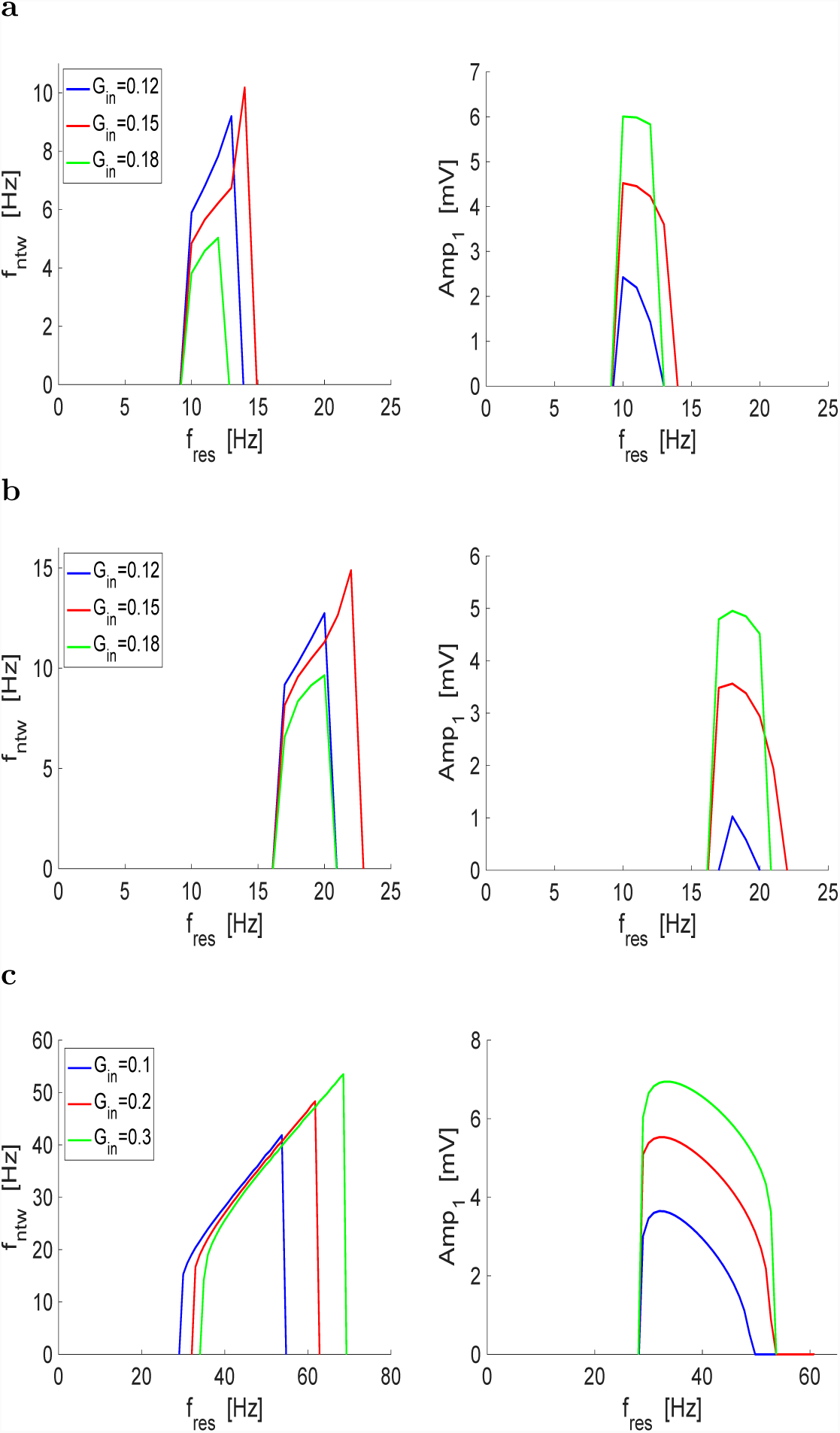
Oscillations in mutually inhibitory resonator-passive cell networks (2D/1D): the intrinsic resonant frequency controls the network frequency. **Left columns.** Network oscillation frequency as a function of *f*_*res*_. **Right columns.** Network oscillation amplitude (oscillator 1) as a function of *f*_*res*_. The synaptic conductances *G*_*in,*1,2_ = *G*_*in,*2,1_ are equal to the values reported in the figure. **a.** *g*_*L,*1_ = 0.25, *Z*_1,*max*_ = 3.9. **b.** *g*_*L,*1_ = 0.25, *Z*_1,*max*_ = 3.7. **c.** *g*_*L,*1_ = 0.1, *Z*_1,*max*_ = 6. We used the following parameter values: *g*_*L,*2_ = 0.5, *E*_*in*_ = −20, *v*_*hlf*_ = 0, *v*_*slp*_ = 1.

The oscillation amplitude increases with increasing values of *G*_*in*_ (= *G*_*in,*1,2_ = *G*_*in,*2,1_) and is more variable than for the self-excited resonator. The oscillatory active *f*_*res*_ band (the range of values of *f*_*res*_ for which network oscillations are possible) is relatively small as compared to the self-excited resonator network and it depends on the value of *Z*_*max*_ and *g*_*L,*1_. All other parameters fixed, decreasing values of *Z*_*max*_ (from Fig. 12-a to -b) causes the oscillatory active resonant frequency band to slide to the right. Fig. 12-c shows that the size active frequency band can be increased by decreasing *g*_*L,*1_ and increasing *Z*_*max*_. A proper comparison would involve changing one parameter at the time, but decreasing values of *g*_*L,*1_ require increasing values of *Z*_*max*_ for the oscillations to be present.

### 3.4 Mutually excitatory 2D/1D hybrid networks can generate sustained (limit cycle) oscillations and their frequency monotonically depends on the intrinsic resonant frequency

In Section 3.2 we showed that self-excited resonators can produce limit cycle oscillations, their frequency monotonically depends on the resonator’s resonant frequency, and relaxation oscillations develop for high enough levels of self excitation. Here we extend our results to include two-cell networks. Because self-excited resonators may be thought of as representing a population of synchronized in phase cells, we expect our results from Section 3.2 to hold of these networks. However, the presence of nonlinearities of cubic type are not apparent from either the model equations or the phase-space diagrams and need to be uncovered using the method developed in Section 3.3.

#### 3.4.1 Oscillations can be generated in 2D/1D hybrid networks and are amplified with increasing levels of mutual excitation

Fig. 14-a1 show the small amplitude oscillations generated in a Hopf bifurcation (Fig. 16) for low enough values of *G*_*ex*_ (*G*_*ex,*1,2_ = *G*_*ex,*2,1_). This oscillations are not identical, because the cells are not identical, but they are synchronized in phase. Fig. 14-b1 shows that increasing values of *G*_*ex*_ lead to oscillations of relaxation type. The dynamic mechanisms of oscillation amplification (Figs. 14-a2 and -b2) as well as the cubic-based mechanisms of generation of relaxation oscillations (Figs. 14-a3 and -b3) are analogous to the mutually inhibitory networks discussed in Section 3.3.

**Figure 13:**
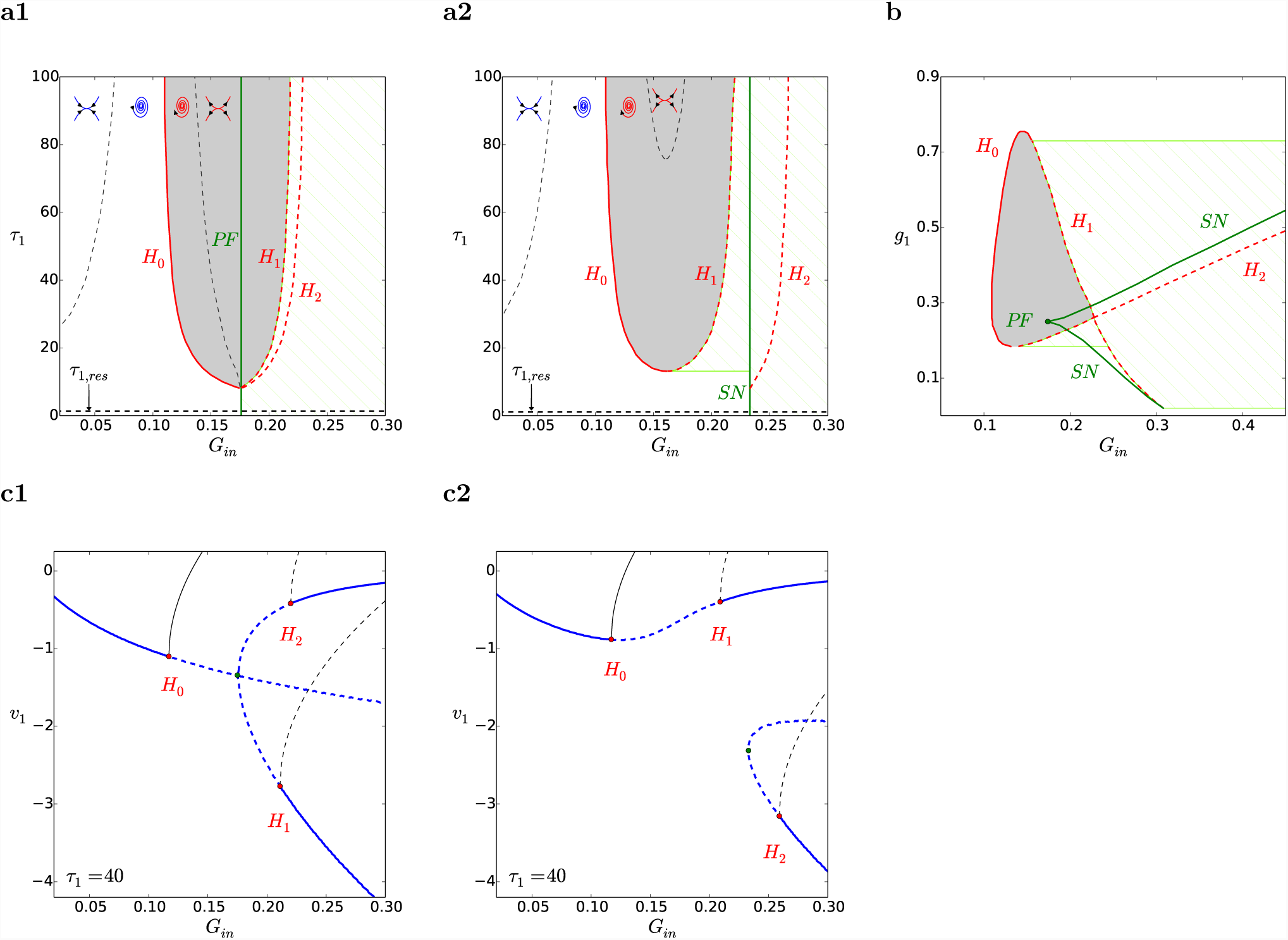
Bifurcation diagrams for mutually inhibitory resonator-passive cell networks (2D/1D) for representative parameter values. The shadowed region corresponds the existence of sustained (limit cycle) oscillations. The green-lined region corresponds to multistability (limit cycle and/or fixed-points). The inset trajectory diagrams indicated the dynamics within the regions bounded by the solid and dashed curves (except the solid green curve): stable nodes, stable foci, unstable foci and unstable nodes (from left to right). The inset diagrams correspond to the 3D linearized system for the fixed-point before the static bifurcation. *H*_0_, *H*_1_ and *H*_2_ note the Hopf bifurcation branches, *PF* notes the pitchfork bifurcation branch and *SN* notes the saddle-node branch. **a.** Bifurcation diagram in *G*_*in*_-*τ*_1_ parameter space. Cell 1 is a resonator for values of *τ*_1_ > *τ*_1,*res*_ (dashed-black horizontal line). **a1** *g*_*L,*1_ = 0.25 and *g*_1_ = 0.25. **a2** *g*_*L,*1_ = 0.25 and *g*_1_ = 0.3. **b.** Bifurcation diagram in *G*_*in*_-*g*_1_ parameter space for *g*_*L,*1_ = 0.25 and *τ*_1_ = 100. **c.** Bifurcation diagram with *G*_*in*_ as bifurcation parameter. The solid- and dashed-blue curves represent stable and unstable fixed-points, respectively. The solid- and dashed-black curves represent the stable and unstable limit cycle branches created at the Hopf bifurcations (red dots). **c1** *g*_*L,*1_ = 0.25, *g*_1_ = 0.25 and *τ*_1_ = 40. **c2** *g*_*L,*1_ = 0.25, *g*_1_ = 0.3 and *τ*_1_ = 40. We used the following parameter values: *g*_*L,*2_ = 0.5, *E*_*in*_ = −20, *v*_*hlf*_ = 0, *v*_*slp*_ = 1.

**Figure 14:**
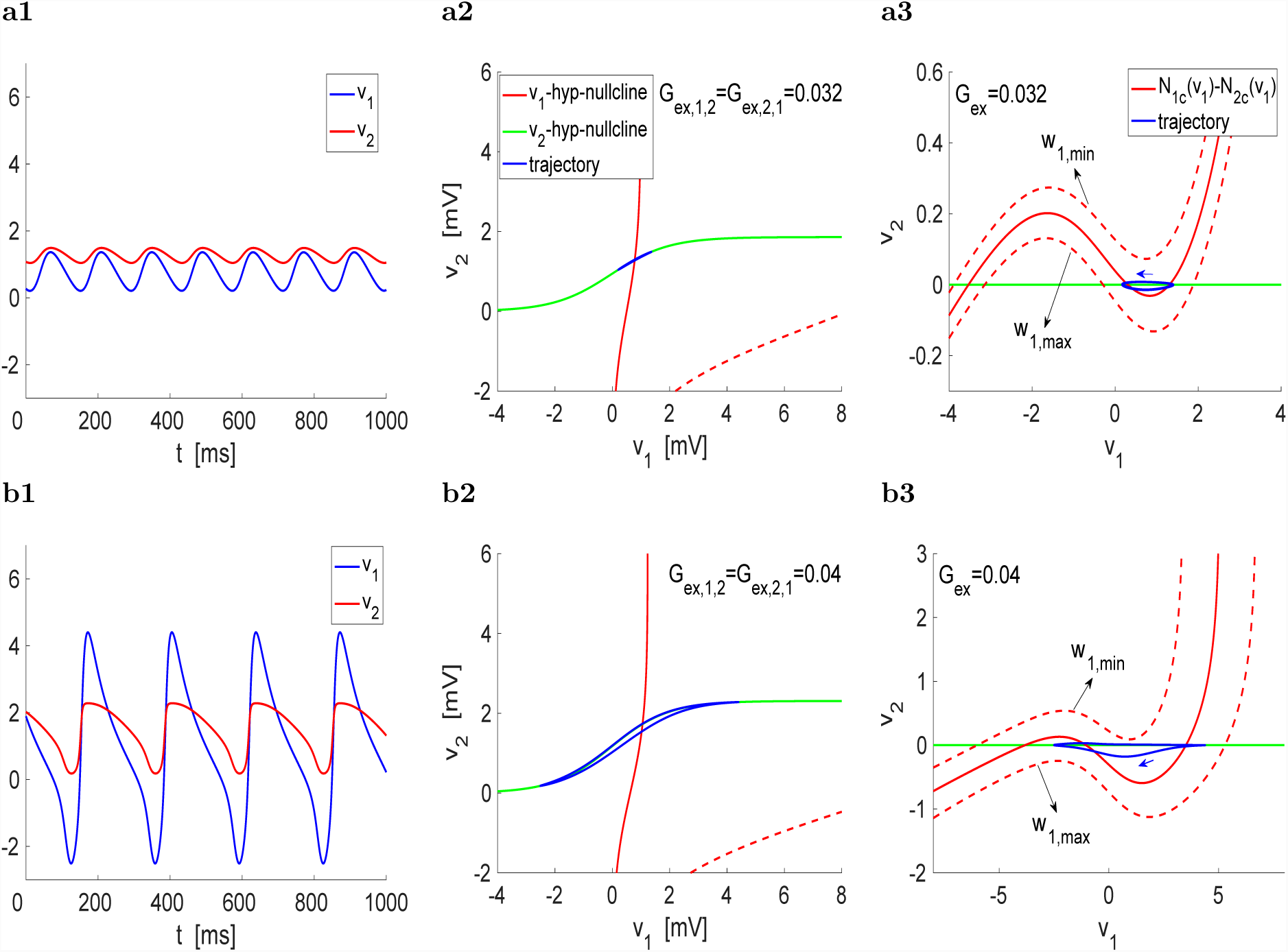
Oscillations generated in mutually excited hybrid 2D-1D networks. Cell 1 is a resonator with *f*_*res*_ = 8 (*f*_*nat*_ = 0) and cell 2 is a passive cell. **Left.** Voltage traces (curves of *v*_1_ and *v*_2_ as a function of *t*). **Middle.** Phase-plane diagrams. The *v*_1_- and *v*_2_-hyper-nullclines are given by (10) and (11), respectively. The dashed red curve represents the *v*_1_ nullcline for cell 1 for *g*_1_ = 0 (no resonant gating variable). **Right.** Adapted phase-plane diagrams relative to the *v*_2_-hyper-nullcline *N*_2*c*_(*v*_1_) (green curve in the phase-plane diagrams in the left panels). The red lines are the differences between the *v*_1_- and *v*_2_-hyper-nullclines in the left panels parametrized by constant values of *w*_1_. The solid-red curve corresponds to an intermediate value of *w*_1_. The dashed-red curves correspond to the maximal *w*_1,*max*_ (lower) and minimal *w*_1,*min*_ (upper) values of *w*_1_. The trajectories (blue curves) are also referred to the *v*_2_-hyper-nullcline *N*2_*c*_(*v*_1_). **a.** *G*_*ex,*1,2_ = *G*_*ex,*2,1_ = 0.032. The network frequency is *f*_*ntw*_ ∼ 7.1. **b.** *G*_*ex,*1,2_ = *G*_*ex,*2,1_ = 0.04. The network frequency is *f*_*ntw*_ ∼ 4.3. We used the following parameter values: *g*_1_ = 1.8, *g*_*L,*1_ = 0.1, *g*_*L,*2_ = 1, *τ* = 750, *E*_*ex*_ = 60, *v*_*hlf*_ = 0, *v*_*slp*_ = 1.

#### 3.4.2 The resonator’s intrinsic frequency controls the network oscillations frequency

Our results are presented in Fig. 15 for (i) values of *Z*_1,*max*_ that increase from panel a to b (for a fixed-value of *g*_*L,*1_), and (ii) values of *g*_*L,*1_ that decrease from panel b to c (for a fixed values of *Z*_1,*max*_). In contrast to the mutually inhibitory networks, by increasing *Z*_1,*max*_, the oscillatory active resonant frequency band is increased, while the onset of oscillations occurs for lower values of *f*_*res*_, similarly to mutually inhibitory networks. The opposite behavior is observed for decreasing *g*_*L,*1_ with all the other parameters fixed. The behavior of the oscillations amplitude is similar to the mutually inhibitory networks.

**Figure 15:**
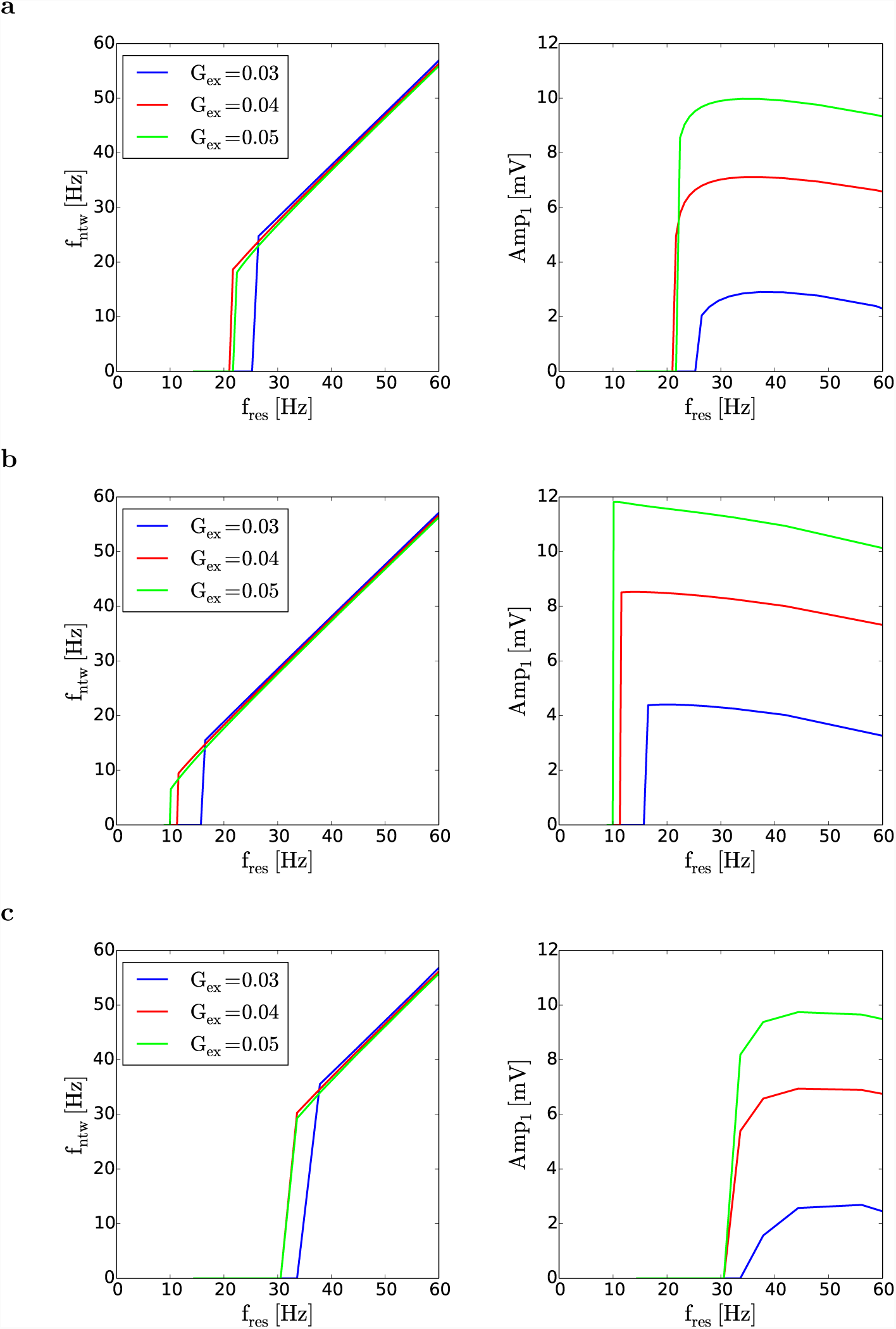
Oscillations in mutually excited hybrid 2D-1D networks: the intrinsic resonant frequency controls the network frequency. **Left columns.** Network oscillation frequency as a function of *f*_*res*_. **Right columns.** Network oscillation amplitude (oscillator 1) as a function of *f*_*res*_. The synaptic conductances *G*_*ex,*1,2_ = *G*_*ex,*2,1_ are equal to the values reported in the figure. **a.** *g*_*L,*1_ = 0.1, *Z*_1,*max*_ = 9.2. **b.** *g*_*L,*1_ = 0.1, *Z*_1,*max*_ = 9.87. **c.** *g*_*L,*1_ = 0.08, *Z*_1,*max*_ = 9.87. We used the parameter value *g*_*L,*2_ = 1, *E*_*ex*_ = 60, *v*_*hlf*_ = 0, *v*_*slp*_ = 1.

**Figure 16:**
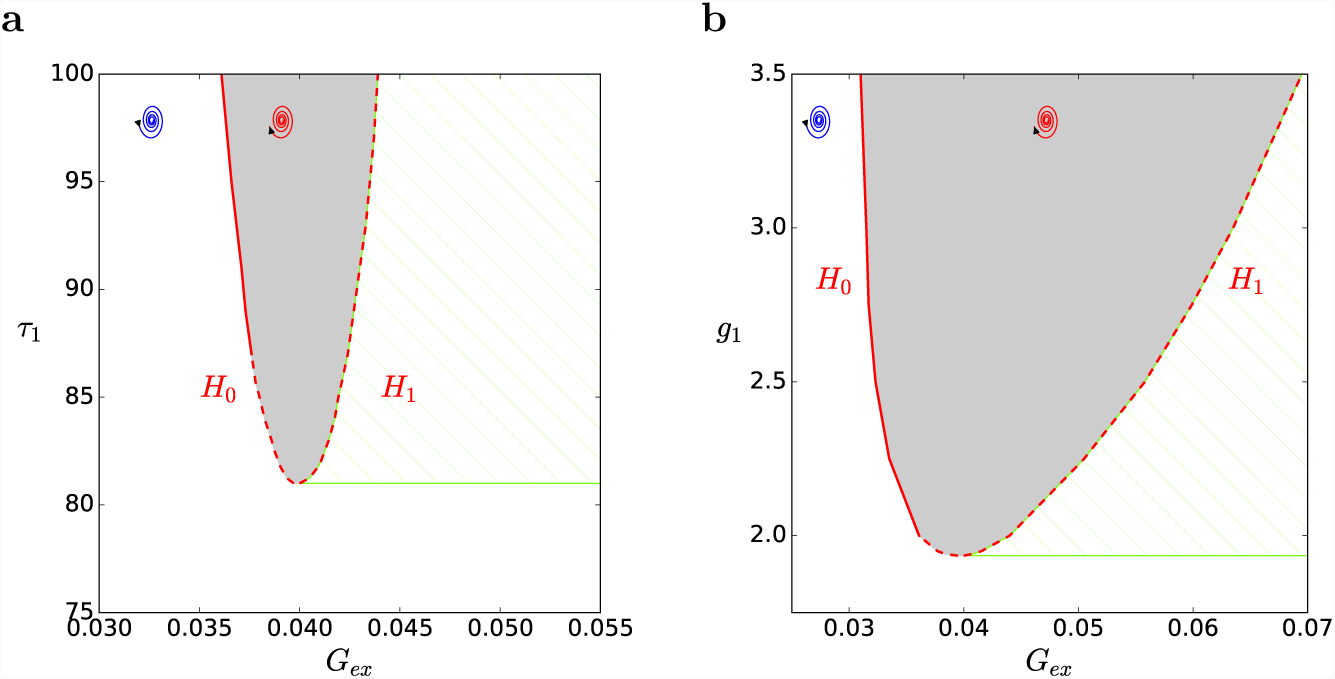
Bifurcation diagrams for mutually excitatory resonator-passive cell networks (2D/1D) for representative parameter values. The shadowed region corresponds the existence of sustained (limit cycle) oscillations. The green-lined region corresponds to multistability (limit cycle and/or fixed-points). The inset trajectory diagrams indicated the dynamics within the regions bounded by the solid and dashed curves (except the solid green curve): stable and unstable foci. *H*_0_ and *H*_1_ note the Hopf bifurcation branches. **a.** Bifurcation diagram in *G*_*ex*_-*τ*_1_ parameter space for *g*_*L,*1_ = 0.1 and *g*_1_ = 2. Cell 1 is a resonator for values of *τ*_1_ > *τ*_1,*res*_ ∼ 0.205: **b.** Bifurcation diagram in *G*_*ex*_-*g*_1_ parameter space for *g*_*L,*1_ = 0.1 and *τ*_1_ = 100. We used the following parameter values: *g*_*L,*2_ = 1.2, *E*_*ex*_ = 60, *v*_*hlf*_ = 0, *v*_*slp*_ = 1.

### 3.5 Graded mutually inhibitory or excitatory 2D/2D resonator networks generate sustained (limit cycle) oscillations and their frequency interact to control the network frequency

Here we extend our results from Sections 3.3 and 3.4 to networks having two mutually connected 2D resonators (that are not damped oscillators). We consider heterogeneous networks of non-identical resonator in order to test the effects of the interaction band-pass filters with different frequency bands. Because the mechanisms of generation of oscillations are similar to those discussed in Sections 3.3 and 3.4, we focus on the effects of the resonant frequencies of the participating resonators on the network oscillation frequency. Our results are presented in Figs. 17. The gray curves corresponds to networks of resonators with the same frequency band. The network model is given by system (1)-(4) with *g*_1_, *g*_2_ > 0.

**Figure 17:**
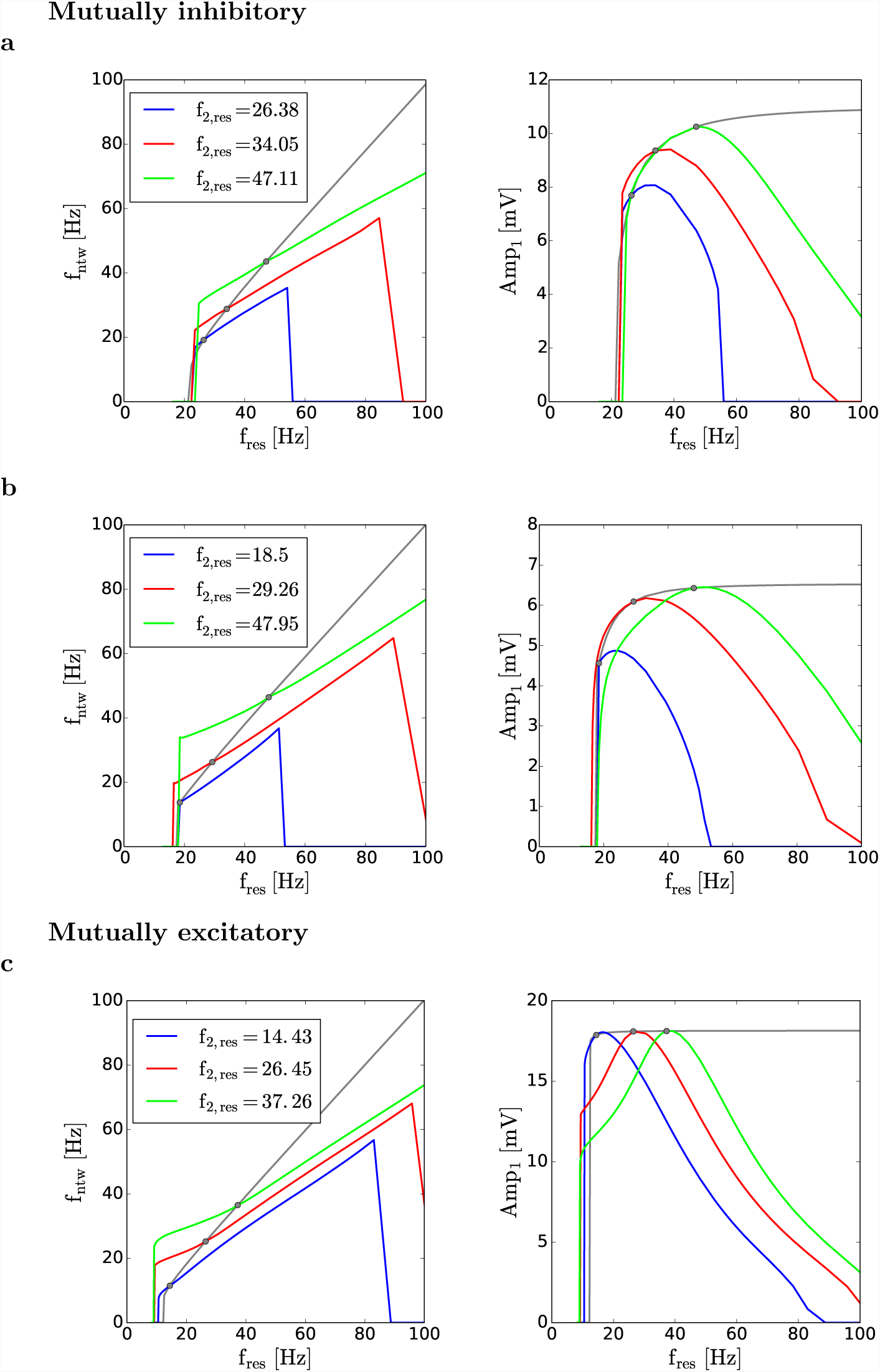
Oscillations in mutually inhibitory or excitatory resonator cell networks (2D-2D): the intrinsic resonant frequencies interact to control the network frequency. **Left columns.** Network oscillation frequency as a function of *f*_*res*_. **Right columns.** Network oscillation amplitude (oscillator 1) as a function of *f*_*res*_. The gray curves correspond to a network of identical cells for fixed values of *g*_*L,*1_ = *g*_*L,*2_ and *Z*_1,*max*_ = *Z*_2,*max*_. The colored curves correspond to a fixed cell 2 with the resonance frequency *f*_2,*res*_ indicated in the figure. **a.** *g*_*L*_ = 0.1, *Z*_*max*_ = 6, *G*_*in*_ = 0.1. **b.** *g*_*L*_ = 0.25, *Z*_*max*_ = 3.7, *G*_*in*_ = 0.1. **c.** *g*_*L*_ = 0.1, *Z*_*max*_ = 9.2, *G*_*ex*_ = 0.03.

Fig. 17 shows the dependence of the network frequency on *f*_1,*res*_ for representative values of *f*_2,*res*_ and other model parameters for mutually inhibitory (panels a and b) and mutually excitatory (panels c) networks. As expected, in all cases the network frequency monotonically depends on the resonant frequency of both oscillators.

The range of values of *f*_1,*res*_ for which network oscillations are possible increase with increasing values of *f*_2,*res*_ as does the network frequency. However, the network frequency and the resonant frequency of the oscillators is no longer one-to-one, as was the case for the 2D/1D networks investigated in Sections 3.3 and 3.4, but it depends on the complex interaction between the two resonators. The one-to-one dependence between the network frequency and the resonant frequencies of the individual oscillators occurs when the two resonators have the same resonant frequency (black dots). The slopes of the network frequency curves for non-identical resonators are smaller than for identical resonators indicating that the network frequency is larger (smaller) than the resonant frequency to the left (right) of the black dot. This is independent of the mechanisms of amplification (mutual inhibition or mutual excitation).

Increasing values of *Z*_2,*max*_ for fixed values of *Z*_1,*max*_ and *f*_2,*res*_ causes the range of values of *f*_1,*res*_ for which oscillations exist increase, but the slope of the network frequency curves remains almost unchanged, independently of the value of *g*_*L,*1_ used and whether the mechanisms of amplification is based on mutual excitation or mutual inhibition (not shown).

### 4 Discussion

Network oscillations emerge from the cooperative activity of the intrinsic properties of the participating neurons and the synaptic connectivity, and involve the nonlinearities and time scales present in the circuit components and these that emerge from their interplay. MPR is a property of the interaction between oscillatory inputs and the intrinsic neuronal properties (intrinsic resonant and amplifying processes) that uncovers a circuit latent time scale associated to the resonant frequency (MPR can be observed in the absence of intrinsic damped oscillations [2, 3]). MPR has been investigated both experimentally and theoretically in many neuron types [1–4, 12–56]. However, whether MPR plays any functional role for network oscillations or is simply an epiphenomenon is largely an open question. A few studies have investigated the oscillatory properties of networks including neurons that exhibit MPR [41, 57–63] or have resonant gating variables [64–66]. But the role that MPR plays in the generation of network oscillations and how the latent time scales affect the properties of the oscillatory networks in which they are embedded remained to be understood. One problem in addressing these questions is the fact that a resonator and its latent oscillatory properties are characterized by the impedance profile and the resonant frequency, respectively, which are properties that only emerge in the presence of an oscillatory (AC) interaction, unlike damped oscillations that are intrinsic neuronal properties that can be uncovered by constant (DC) perturbations.

In this paper we set out to investigate these issues using minimal network models consisting of non-oscillatory resonators mutually coupled to either a low-pass filter neuron or another band-pass filter (resonator). In this way we could separate the different effects that give rise to network oscillations in two different levels of organization that can be manipulated separately. The resonator provides the negative feedback and the network connectivity provides the amplification. Because we leave out resonators that can be also damped-oscillators, the network oscillations are not inherited from the individual cell level, but are created by the combination of the individual cell and connectivity properties.

We showed that oscillations can be generated in networks of increasing complexity: (i) self-excited band-pass filters, (ii) mutually inhibited band- and low-pass filters, (iii) mutually excited band- and low-pass filters, (iv) mutually inhibited band-pass filters, and (v) mutually excited band-pass filters. The presence of a resonator is necessary to generate oscillations in these networks; if the resonators are substituted by low-pass filters, network oscillations are not possible. However, what characterizes the oscillatory activity of a resonator is the resonant frequency, which cannot be assessed in the absence of oscillatory inputs. By showing that the network frequency monotonically depends on the resonant frequency of the individual band-pass filters, we provide a direct link between MPR and the generation of network oscillations. To our knowledge, this is the first time such a link is provided. A similar results was obtained in electrically coupled networks [41], but in these cases, the network oscillations were driven by one of the nodes that was a sustained oscillator. Network oscillations have been shown to emerge as the result of the interaction of damped oscillators [65, 66, 82], but in these cases, the network oscillations are inherited from the oscillatory activity of the individual intrinsically oscillatory nodes.

The existence of sustained oscillations in networks of non-oscillatory neurons is not without precedent. The inferior olive oscillatory network studied in [65] is composed of electrically coupled neurons that, when isolated, are damped oscillators. In this case, the individual neurons are nonlinear and include both resonant and amplifying effects, but the connectivity is linear. The model investigated in [64] involves nonlinear neurons reciprocally inhibited with graded synapses. For baseline values of the DC input current, the neurons are quasi-linear and are at most damped oscillators. The nonlinearities developed for negative input current values combined with the dynamics resulting from the mutual synaptic inhibition result in the post-inhibitory rebound mechanism underlying the observed network oscillations. Post-inhibitory rebound (PIR) and subthreshold resonance are closely related phenomena since both require the presence of a negative feedback effect, but they are different in nature. The mechanisms investigated in [64] depend crucially on the effective pulsatile nature resulting from the dynamic interaction between cells and synaptic connectivity. Models having an h-current also show PIR. Even in the presence of an (additive) amplifying current, such as the *I*_*h*_ + *I*_*Nap*_ model, the functional connectivity in these models is not PIR-based, but rather resonance-based as we show in this paper (unpublished observation). In [67, 73] oscillations emerge in two reciprocally inhibited passive cells where one of them is self-excited, thus providing additional dynamics to the network. The model studied in [41] consists of an oscillator electrically coupled to a follower resonator whose intrinsic resonant frequency directly affects the network frequency while the shape of the impedance profile remains almost unchanged.

The minimal models we used in this paper serve the purpose of establishing the role of MPR for the generation of network oscillations. Other types of models could include resonant properties at the network level. Moreover, there are alternative possible scenarios where, for example, amplification occurs at the single cell level and the negative feedback effect occurs at the network level. These types of networks are beyond the scope of this paper. The understanding of the oscillatory properties of such networks requires more research.

The types of models we used could be argued to be too simplistic and not realistic. We used these models precisely because of their simplicity in order to understand some conceptual points that can be generalized and applied to more realistic networks. However, one should note that the type of models we used are very close to the firing rate models of Wilson-Cowan type [69] with adaptation [70–72], which are essentially resonators (unpublished observation). In this models, the nonlinearity is similar to the one we used (sigmoid type, instantaneously fast). Therefore, our results can be easily generalized to these models. One example are the networks of OLM cells and fast spiking (PV^+^) interneurons (INT) that have been shown to be able to produce network oscillations [83, 84]. OLM cells show MPR [33], while the presence of MPR in INT is debated [25, 33]. Although our models are simplistic, they make predictions that can be tested using the dynamic clamp technique [85, 86].

Our results open several questions regarding the ability of networks of band-pass filters to generate oscillatory patterns and how the properties of these patterns depend on the properties of the band-pass filters. More research is required to address these issues.

## Acknowledgments

This work was partially supported by the National Science Foundation grant DMS-1608077 (HGR) and the Universidad Nacional del Sur grant PGI 24/L096 (AB). The authors thank Eran Stark for useful comments and discussions. HGR is grateful to the Courant Institute of Mathematical Sciences at NYU and the Department of Mathematics at Universidad Nacional del Sur, Argentina.

## A Graded networks of passive cells: linearization

The linearization of system (1) with *g*_*k*_ = 0 for *k* = 1, 2 and *I*_*syn,k*_ given by (3) and (4) reads

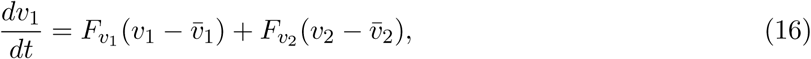

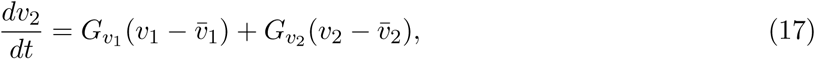

where

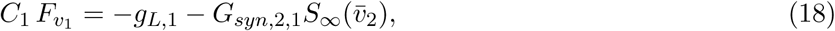

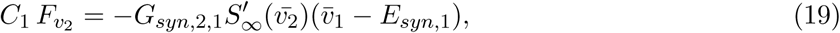

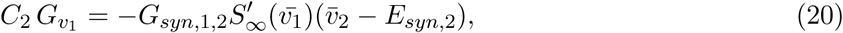

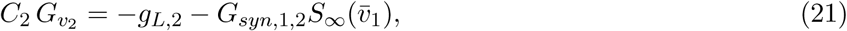

The eigenvalues (*r*_1_ and *r*_2_) are given by

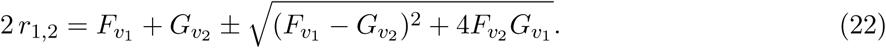

The first two terms in (22) are always negative (provided *g*_*L,*1_ > 0 and *g*_*L,*2_ > 0). The second term in the radicand is positive if *F*_*v*_2 and *G*_*v*_1 have the same sign and negative if *F*_*v*_2 and *G*_*v*_1 have different signs. Therefore, the fixed-point for networks with the same type of connections (both excitatory or both inhibitory) can be either stable nodes or saddles, while the fixed-points for excitatory-inhibitory networks can be either stable nodes or stable foci.

## B Impedance profiles with fixed peak values and changing resonant frequencies

The impedance profile for a 2D system of the form

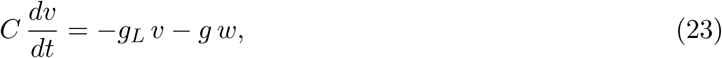

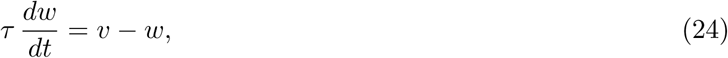

is given by

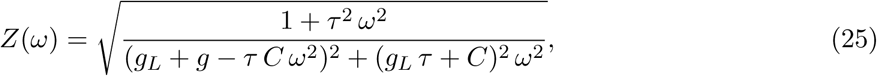

where *ω* = 2*πf/*1000. The resonant frequency is given by

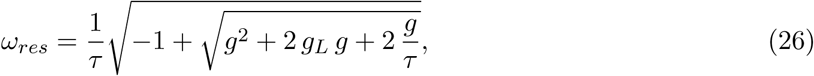

where for simplicity *C* = 1. The impedance peak is given by

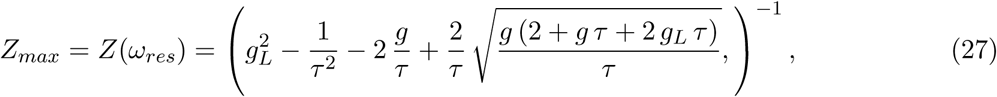

from where

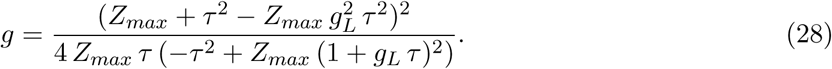

In order to calculate the values of *g* and *τ*, if they exist, for given values of *Z*_*max*_ and *g*_*L*_ (fixed) we proceed as follows. First we take values of *τ* within certain range and compute the corresponding values of *g* using (28). For these values of *g*_*L*_, *g* and *τ* we compute *ω*_*res*_ using (26) and *C* = 1. In this way we have *g* = *g*(*τ*) and *ω*_*res*_ = *ω*_*res*_(*τ*) for a given value of *Z*_*max*_.

